# Spatially pooling photon information enables photon-efficient quantitative imaging

**DOI:** 10.64898/2026.08.19.745572

**Authors:** Wonsang Hwang, Iván Coto Hernández, Conor L. Evans

**Affiliations:** Wellman Center for Photomedicine, Massachusetts General Hospital, Charlestown, MA, USA; Center for Interdisciplinary Innovation in Imaging, Massachusetts General Hospital, Charlestown, MA, USA

**Keywords:** fluorescence lifetime imaging, hyperspectral imaging, Poisson inverse problems, photon efficiency, Cramér–Rao bound

## Abstract

Quantitative fluorescence imaging techniques such as fluorescence lifetime imaging microscopy and hyperspectral imaging infer molecular contrast from photons distributed across spatial pixels and temporal or spectral channels. In the few-photon regime, however, conventional pixel-wise analysis discards the spatial relationships imposed across neighboring pixels by the microscope point-spread function (PSF). Here we show that this spatially distributed information can be recovered without prior knowledge of emitter positions, spatial support or component assignments. We introduce SPOOL (Spatially Pooled Optical Observation Likelihood), a training-free Poisson inverse framework that jointly recovers source-space amplitudes and quantitative contrast by combining the PSF with temporal-decay or spectral-response dictionaries. For an isolated source, the attainable precision gain is governed by a dimensionless optical quantity: the PSF width expressed in detector pixels. The predicted gain therefore scales with optical sampling rather than with the physical origin of the contrast. The model predicts that lifetime-precision gain scales approximately linearly with the number of pixels spanning the PSF full width at half maximum, a scaling reproduced by Monte Carlo simulations. At one detected photon per foreground pixel, the reconstruction reduces lifetime dispersion sixfold in fluorescent-bead experiments and decreases the lifetime root-mean-square error relative to a high-photon reference from 1.19 to 0.45 ns in dual-labeled cells. The same framework extends directly to hyperspectral imaging, recovering spectral contrast from generic emission bands without prior fluorophore spectra.

## 1 Introduction

Quantitative fluorescence imaging becomes unreliable in the few-photon regime, where shot noise and background destabilize pixel-level parameter estimation [1–4]. This limitation is particularly acute in fluorescence lifetime imaging microscopy (FLIM) and hyperspectral imaging because the detected photon budget is further partitioned across temporal or spectral channels before a molecular parameter can be estimated [3, 5, 6]. Increasing the acquisition time or excitation intensity can raise the detected photon count, but longer exposures sacrifice imaging speed, whereas higher light dose accelerates photobleaching and increases the risk of phototoxicity [7–9]. These trade-offs are especially restrictive in fast live-cell imaging, deep-tissue microscopy and single-molecule measurements, where sample dynamics, optical scattering and finite fluorophore photon budgets constrain acquisition [7, 10, 11].

Photon scarcity has therefore usually been treated as a problem of extracting a more reliable estimate from each pixel’s sparse decay curve or spectrum. FLIM commonly uses least-squares or maximum-likelihood fitting and fit-free phasor analysis [12–14], whereas hyperspectral data are typically analyzed using non-negative linear unmixing or spectral phasor analysis [15, 16]. More recent methods use denoising, direct parameter estimation and learned spatial context to recover useful contrast from increasingly sparse measurements [4, 17–19]. Noise-aware spectral inference has likewise improved unmixing under low-signal conditions [1, 3], while physics-informed and self-supervised reconstruction methods have improved fidelity and robustness by incorporating image formation into inference [20, 21]. These approaches improve how noisy measurements are interpreted, but leave a more fundamental question unresolved: whether all of the information recorded by the optical system has entered the estimation problem in the first place.

A conventional diffraction-limited microscope distributes the detection probability of photons from each source across neighboring image pixels according to the point spread function (PSF) [22, 23]. Photons detected at neighboring pixels may therefore share common source contributions and report on the same lifetime or spectral composition. Conventional FLIM and hyperspectral estimators nevertheless treat the decay curve or spectrum recorded at each detector pixel as a separate inference problem [13, 14, 16]. For an isolated source whose position and support are known, photons distributed across its PSF footprint can be pooled into a single estimate. Real images present a harder problem: source positions and boundaries are unknown, neighboring structures overlap, and different sources may carry distinct temporal or spectral signatures. Structured detection addresses a related axial assignment problem by adding detector-space diversity [24]; conventional image-plane measurements provide no equivalent cue for lateral source assignment. The relevant challenge is therefore not merely to pool photons, but to recover their shared source information without knowing their assignments in advance.

Previous computational microscopy approaches have shown that information distributed across temporal, detector or angular coordinates can be recovered by explicitly modelling multidimensional image formation [21, 24–26]. Multi-image deconvolution has also been applied to time-resolved data to improve spatial resolution and signal-to-noise ratio, but did not retain the temporal dimension as an explicit variable for quantitative lifetime estimation [27]. Here, we address the complementary problem of using the known spatial redistribution of photons to improve estimation along a temporal or spectral contrast axis. In this view, the PSF is not merely a source of blur, but a known forward-model relationship linking measurements that report on the same underlying source. We show that preserving this information requires the spatial and contrast dimensions to remain coupled during inference: sequential pixel-wise decomposition and deconvolution cannot fully restore the spatial–contrast relationships discarded during the initial estimation step.

We therefore formulate few-photon quantitative imaging as a joint spatial–contrast inverse problem. As illustrated in Fig. 1, the forward model represents the expected photon-count cube as a sum of non-negative source-space component maps, each convolved with the microscope PSF and weighted by a basis profile along the contrast axis, with photon detection described by Poisson statistics [28–30]. Non-negative basis representations are widely used for quantitative unmixing [1, 3, 15]; here, however, the spatial and contrast dimensions are coupled within a single likelihood. The resulting training-free estimator, SPOOL (Spatially Pooled Optical Observation Likelihood), jointly recovers component amplitudes and their associated quantitative contrast directly from the raw photon counts, allowing photons detected across neighboring pixels to constrain the same source location while resolving overlapping structures with distinct temporal or spectral signatures. The same formulation also predicts the attainable precision: for an isolated source, the gain over single-pixel estimation is determined by the PSF width expressed in detector pixels, independent of whether the contrast is temporal or spectral. We show that SPOOL recovers most of this available precision in the few-photon regime, outperforms sequential unmixing followed by deconvolution, and extends directly from FLIM to hyperspectral imaging. To facilitate broader use and reproducibility, we provide SPOOL as an open-source, user-friendly plugin for ImageJ. Together, these results demonstrate that quantitative information redistributed by the optics can remain recoverable when conventional pixel-wise inference fails.

**Fig. 1.**
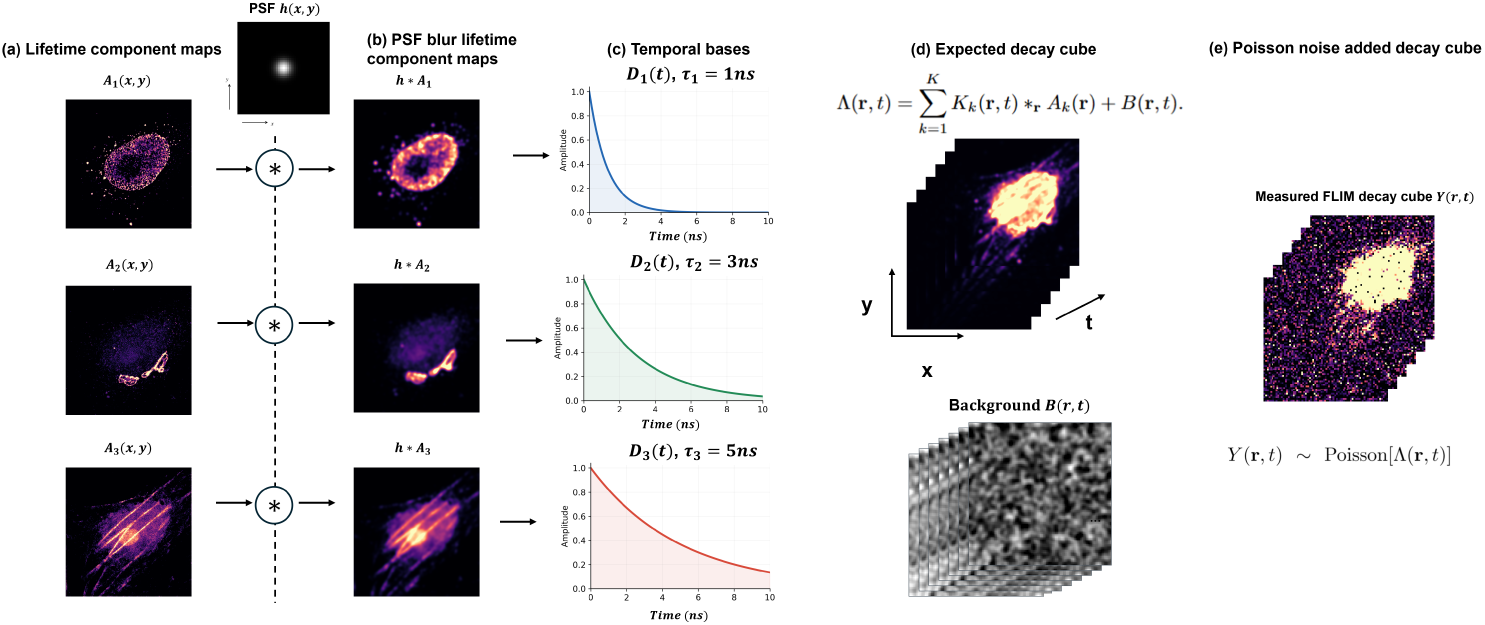
Photons carrying the same molecular information are spatially spread by the optics and then analyzed separately. The forward model makes both facts explicit. **a**, The specimen is represented by nonnegative source-space component amplitude maps *A_k_*(*x, y*), one per basis component. **b**, Each map is spatially blurred by the same point spread function *h*(*x, y*): photons emitted at one location are distributed over a neighbourhood of detector pixels. **c**, Each component is weighted by its own basis profile *D_k_*(*t*) along the contrast axis – a temporal decay basis for FLIM, a spectral emission band for hyperspectral imaging. **d**, The expected photon distribution Λ(*x, y, t*) is the sum of the blurred, basis-weighted components plus a background term *B*(*x, y, t*). **e**, The measurement is a Poisson realization of that distribution, *Y* (*x, y, t*) *∼* Poisson(Λ(*x, y, t*)). A pixel-wise estimator reads the decay histogram at each detector pixel in isolation, discarding the relationship that **b** imposed between them. The inverse problem instead recovers *{A_k_}* from *Y* by maximizing the Poisson likelihood of the raw counts under the same forward model, so that photons distributed away from a source jointly constrain its recovered source-space components.

## Methods

### Spatial–contrast Poisson reconstruction

Let *Y* (**r***, t*) denote the measured photon count at spatial position **r** = (*x, y*) and contrast-axis bin *t*. The sample is represented by *K* nonnegative source-space component maps *A_k_*(**r**), each associated with a unit-area basis profile *D_k_*(*t*). For FLIM, *D_k_*(*t*) is an instrument-response function (IRF) convolved with an exponential decay; for hyperspectral imaging, it is a normalized Gaussian spectral band. The expected photon-count distribution is

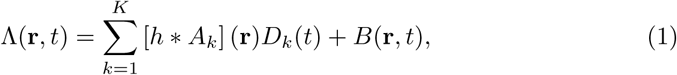

where *h* is the unit-sum spatial point spread function (PSF), denotes twodimensional convolution over **r**, and *B*(**r***, t*) represents background photon contributions from sources such as detector dark counts or autofluorescence. Photon detection is modeled as *Y* (**r***, t*) ~ Poisson[Λ(**r***, t*)]. The amplitude maps are estimated by minimizing

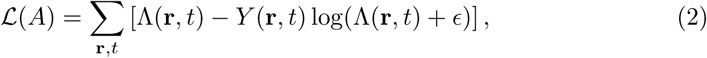

subject to *A_k_*(**r**) 0.

We use a damped multiplicative update related to Richardson–Lucy and expectation–maximization reconstruction [28–30]. At iteration *m*,

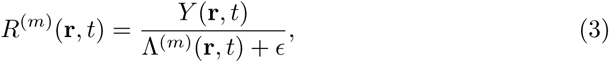

is the ratio of observed to predicted counts. This is a multiplicative Poisson residual that exceeds unity where the current estimate underpredicts the data and falls below unity where it overpredicts. This residual is projected back to source space for each component,

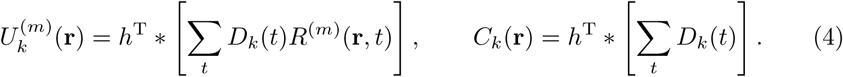

The estimate is then updated as

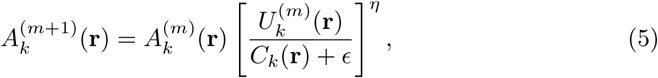

where 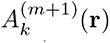 denotes the updated estimate of the *k*-th source-space component map at iteration *m* + 1, *η* ∈ (0, 1] is a damping exponent that shortens each multiplicative step (with *η* = 1 corresponding to the undamped Richardson–Lucy update), the sum over *t* correlates the residual with the basis profile *D_k_*, and the adjoint PSF *h*^T^(**r**) = *h*(*−***r**) redistributes the result from detector space to source space. The normalization *C_k_* is the sensitivity image, the response of the same back-projection to uniform data, so that the ratio 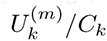 equals unity at convergence; it departs from unity only near the field boundary, where part of the PSF footprint falls outside the measured region.

All SPOOL reconstructions used 50 iterations and *η* = 0.9, fixed across every dataset rather than tuned per sample. Multiplicative Richardson–Lucy-type updates are semi-convergent: additional iterations initially reduce deterministic reconstruction bias, but continued iteration progressively amplifies shot noise, so the iteration count acts as a bias–variance operating point rather than a convergence criterion [31]. We therefore selected a common iteration count from the ground-truth single-emitter simulation rather than tuning it separately for each dataset or photon level. The RMSE-minimizing iteration count shifted with photon level, with lower-count data favoring earlier stopping and higher-count data tolerating additional iterations. We therefore averaged the lifetime RMSE over the complete 1–15 photons-per-pixel range. The resulting curve showed a broad, shallow minimum around 50–75 iterations. Repeating the analysis with the PSF width supplied to the reconstruction perturbed by ±10% produced minima in the same range, with 50 iterations lying near the minimum for all three PSF assumptions. We consequently fixed the reconstruction at 50 iterations for every simulated and experimental dataset, including the hyperspectral reconstruction. The exponent *η* damps each multiplicative step, with *η* = 1 corresponding to the undamped Richardson–Lucy update and *η <* 1 shortening the step along the same ascent direction. We fixed *η* = 0.9 throughout rather than tuning it for individual datasets; sweeping *η* from 0.7 to 1.0 produced less than a 1% variation in reconstruction error, indicating little sensitivity to the damping parameter over this range.

The background *B* was held fixed rather than jointly estimated, taken as spatially and temporally uniform, and estimated for each dataset from a signal-free corner of the field; the same background model and value were supplied to every method compared. Initialization, background estimation, FFT boundary handling, numerical safeguards, and dictionary construction are described in Supplementary Methods S1 and S2.

Setting *h* to a Dirac delta function removes PSF-mediated coupling between neighboring object locations, such that the likelihood factorizes spatially and each detector pixel can be estimated independently. This limiting case corresponds to pixel-wise Poisson maximum-likelihood estimation and serves as our primary training-free baseline. Importantly, the Dirac-delta assumption is applied only to the reconstruction model and does not imply that the acquired measurements are free of optical blur. Rather, the pixel-wise estimator treats the PSF-convolved detector-space amplitudes, *A_k_*, as independent unknowns and therefore does not exploit the spatial information encoded by the optical PSF.

For SPOOL, the recovered source-space quantitative parameter map is computed from the component amplitudes as

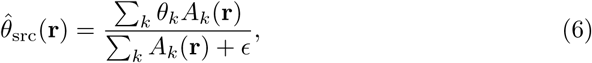

where *θ_k_* = *τ_k_* for FLIM and *θ_k_* = *λ_k_* for hyperspectral imaging. For the pixel-wise baseline, the same expression is evaluated using the detector-space component maps *A_k_* = *h ∗ A_k_*, which are treated as independent unknowns.

### Optics-derived precision gain

We quantify the theoretical precision gain using the Fisher information and the associated Cramér–Rao lower bound, following established formulations for parameter estimation in photon-limited fluorescence microscopy [23, 32, 33].

For an isolated emitter carrying a contrast parameter *θ* (the lifetime *τ* for FLIM) at **r**_0_, the forward model reduces to

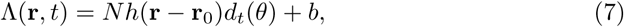

where *N* is the total signal photon count, *d_t_*(*θ*) is the unit-area contrast-axis profile, and *b* is the background. The per-photon Fisher information about *θ*, evaluated for the background-free profile, is

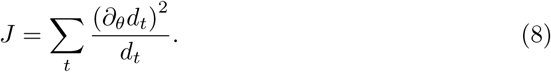

Let *h*_0_ denote the fraction of an emitter’s photons detected in its central pixel. An estimator using only that pixel has Fisher information *Nh*_0_*J*, whereas an estimator using the full PSF footprint has, in the background-free limit,

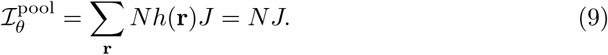

The corresponding gain in standard deviation is therefore

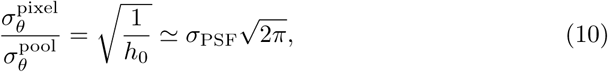

where the final expression applies to a finely sampled, normalized two-dimensional Gaussian PSF and *σ*_PSF_ is expressed in detector pixels. The contrast-axis information *J* sets the absolute precision but cancels from the gain ratio.

This gain is defined relative to estimating the source parameter from the decay recorded at its central detector pixel, as conventional pixel-wise mapping does. It is not a claim that a full-field pixel-wise map contains no further information when the source support is known and the per-pixel estimates are subsequently pooled; that operation is what the oracle estimator performs, and it is included here as a control.

Equation (10) describes an isolated emitter with known position and support and is used as an oracle information reference, not as the exact Cramér–Rao bound of the full-field reconstruction. Numerical bounds include the measured time window, instrument response, background, and unknown amplitude. Further details are provided in Supplementary Methods S3.

For all datasets, we used a theoretical Gaussian PSF with a nominal full width at half maximum of 200 nm. For the single-photon-excitation datasets, this corresponds to the diffraction-limited lateral resolution 0.51*λ/*NA for *λ* = 550 nm and NA = 1.40; the two-photon bead dataset gives a comparable value of 196 nm from the corresponding two-photon excitation model. The PSF was integrated over each detector pixel and converted to detector-pixel units using each dataset’s pixel pitch. The pitch differs between datasets: 30 nm for the beads, 39 nm for the microtubules, 50 nm for the hyperspectral cell and for all simulations, and 60.4 nm for the dual-labeled FLIM cell, corresponding to 6.7, 5.1, 4.0 and 3.3 detector pixels per full width at half maximum. The predicted gain therefore differs between datasets, and each is compared against its own prediction. The PSF width was fixed by this rule and was not fitted to reconstruction performance, with further details provided in the Supplementary Information and Supplementary Tables S1 and S2; the consequences of PSF misspecification are quantified in the Supplementary Information.

### Simulation and experimental validation

The single-emitter simulation used 400 independent Poisson trials at each photon level and compared four estimators: converged pixel-wise Poisson maximum-likelihood estimation (MLE) [13], 3×3-binned MLE, an oracle region-of-interest (ROI)-pooled MLE given the true emitter position and support, and SPOOL. Here, “oracle” denotes an estimator given privileged access to the ground-truth emitter position and spatial support, information that would not be available in a practical experiment. The binning grid was aligned so that the emitter fell at the center of its bin, whereas the oracle pooled the decay histograms over a circular ROI of radius three PSF standard deviations centered on the true emitter position and fitted a single continuous lifetime by Poisson maximum likelihood, without use of the discrete lifetime dictionary. The oracle therefore provides a reference for the performance attainable when the emitter support is known exactly.

Multi-emitter simulations contained homogeneous (*τ* = 2 ns) or heterogeneous (*τ* = 2 and 4 ns) bead phantoms; five methods were compared on eight independent scenes at each of five photon budgets. The true lifetimes lie midway between reconstruction-dictionary nodes, the least favourable position for grid quantization. Both phantoms used the pixel-integrated Gaussian PSF with a 200 nm full width at half maximum and 50 nm pixel sampling of the single-emitter study, allowing direct comparison between the two analyses, and the background level was scaled proportionally with the foreground signal at each photon level, maintaining a mean foreground-to-background ratio of 50. All dictionary-based methods — pixelwise MLE, NC-PCA followed by unmixing, SPOOL and the sequential controls — were reconstructed with the same lifetime-agnostic *K* = 11 dictionary used on the experimental data, so that none was given the true lifetimes.

Experimental validation of SPOOL was carried out through direct comparisons with state-of-the-art computational methods, including (1) pixel-wise MLE [13] and (2) noise-corrected principal component analysis (NC-PCA) [34] followed by nonnegative least-squares unmixing [15]; per-pixel phasor analysis [14], applied without spatial binning or filtering so that its spatial sampling matched the other estimators; the few-photon fluorescence lifetime imaging (FPFLI) network [4], whose local lifetime estimator was retrained on data drawn from this forward model using the authors’ training script and hyperparameters (Supplementary Methods S4); and sequential unmixing–deconvolution controls, in which each detector pixel was first unmixed with the same lifetime dictionary and the resulting amplitude maps were then deconvolved by Wiener, total-variation or Richardson–Lucy reconstruction [28, 29]. The Richardson–Lucy control used the same PSF, multiplicative update and iteration count as SPOOL; it differs from it in the ordering of the two operations and in the least-squares noise model of its unmixing stage. Complete simulation parameters, baseline implementations and hyperparameters are provided in Supplementary Methods S3 and S4.

SPOOL and the above-mentioned methods were compared using fluorescent beads, immunolabeled microtubules, dual-labeled FLIM reference cells, and a dual-labeled hyperspectral cell. Sample preparation, acquisition settings and all reconstruction parameters are described in greater detail in Supplementary Methods S5 and Supplementary Tables S1 and S2.

All simulations, baseline implementations and reconstructions were written in Python. The forward model, Poisson sampling and iterative reconstructions were implemented in PyTorch and accelerated through parallelization on a graphics processing unit (GPU; NVIDIA GeForce RTX 3060 Ti), with the contrast-axis contraction expressed as a matrix product and the spatial convolution as a zero-padded fast Fourier transform. NC-PCA, nonnegative least-squares unmixing and phasor analysis were run on the CPU using NumPy and SciPy. The dictionary profiles and all Cramér– Rao bounds were evaluated in double precision. At the fixed 50-iteration setting with *K* = 11, all experimental datasets reconstructed in under 2 s using double-precision GPU computation and under 0.35 s using single precision, with peak GPU memory usage below 1.9 GiB (see Supplementary Information).

### Photon thinning and evaluation

Experimental photon-budget sweeps were generated by randomly subsampling the detected photons in each raw photon-count cube using binomial thinning. Each recorded photon was retained with probability *p* = *n/n*_full_, and the background was scaled by the same factor. Six independent thinnings were generated at each photon level. Photon budgets are quoted throughout, in simulation and in experiment, as the mean number of detected photons over pixels receiving at least 15% of the peak intensity.

Two references were used. In simulation, accuracy was evaluated against the ground-truth lifetime or wavelength, known by construction. For the experimental samples, each thinned reconstruction was compared with the same method’s own full-photon reconstruction; these values are reported as reference-relative throughout.

Let *θ̂_i_*(**r**) denote the parameter recovered at pixel **r** in thinning *i*, and *θ*^ref^ (**r**) the corresponding reference value – the ground truth in simulation, or that method’s full-photon estimate in experiment. The signed error is 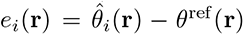, and the root-mean-square error over a mask *M* of *N_M_* pixels and *T* thinnings is

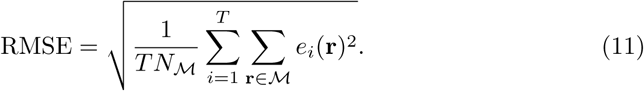

This decomposes as RMSE^2^ = bias^2^ + SD^2^, where the bias is the mean signed error ē and the standard deviation is the dispersion of *e_i_*(**r**) about ē. We report the two terms separately because the estimators compared here differ in bias as well as in dispersion, and a standard-deviation summary alone would conceal a systematic offset from the reference. Reference-relative standard deviation is reported for the single-species samples, in which the mean signed error was negligible, and reference-relative RMSE wherever it was not. In the multi-emitter simulations, errors were evaluated at emitter centers, where the ground-truth lifetime of the contributing source is unambiguous.

Each method was evaluated over its own recovered support, defined by an amplitude threshold of 5% of its maximum for FLIM and 3% for hyperspectral imaging, so that no method is scored on pixels it did not assign signal to. Full metric definitions, mask rules and power-law fitting procedures are provided in Supplementary Methods S6.

## 2 Results

### 2.1 Joint inference recovers information distributed by the point spread function

The PSF distributes photons from a point emitter across neighboring detector pixels, but conventional pixel-wise lifetime estimation analyses each detected histogram independently. SPOOL instead fits the raw photon-count cube under the spatial forward model, allowing photons across the PSF footprint to constrain the same source lifetime. For a normalized two-dimensional Gaussia*_√_*n PSF, the predicted gain in standard deviation over single-pixel estimation is 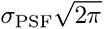 (Eq. (10)). For a 200 nm PSF sampled at 50 nm — four pixels per full width at half maximum — this corresponds to a 4.3*×* gain.

We tested this prediction using a single emitter of known lifetime *τ*_0_ = 2.0 ns at the center of a 48×48-pixel field (2.40×2.40 µm at 50 nm sampling), with 400 independent Poisson trials at each photon level. The foreground comprised the 37 pixels receiving at least 15% of the peak photon rate. This ratio is fixed by the PSF geometry: the central pixel receives 2.3× the foreground mean, so a mean of one photon per pixel places 2.3 photons in the central pixel and 43 photons across the emitter’s full footprint. The single-pixel and pooled bounds are evaluated at those two counts, respectively. At this budget the pixel-wise estimate is dominated by shot noise, whereas the joint reconstruction returns a compact source-space estimate at the original sampling; 3×3 binning reduces the scatter but coarsens the spatial sampling (Fig. 2a).

**Fig. 2.**
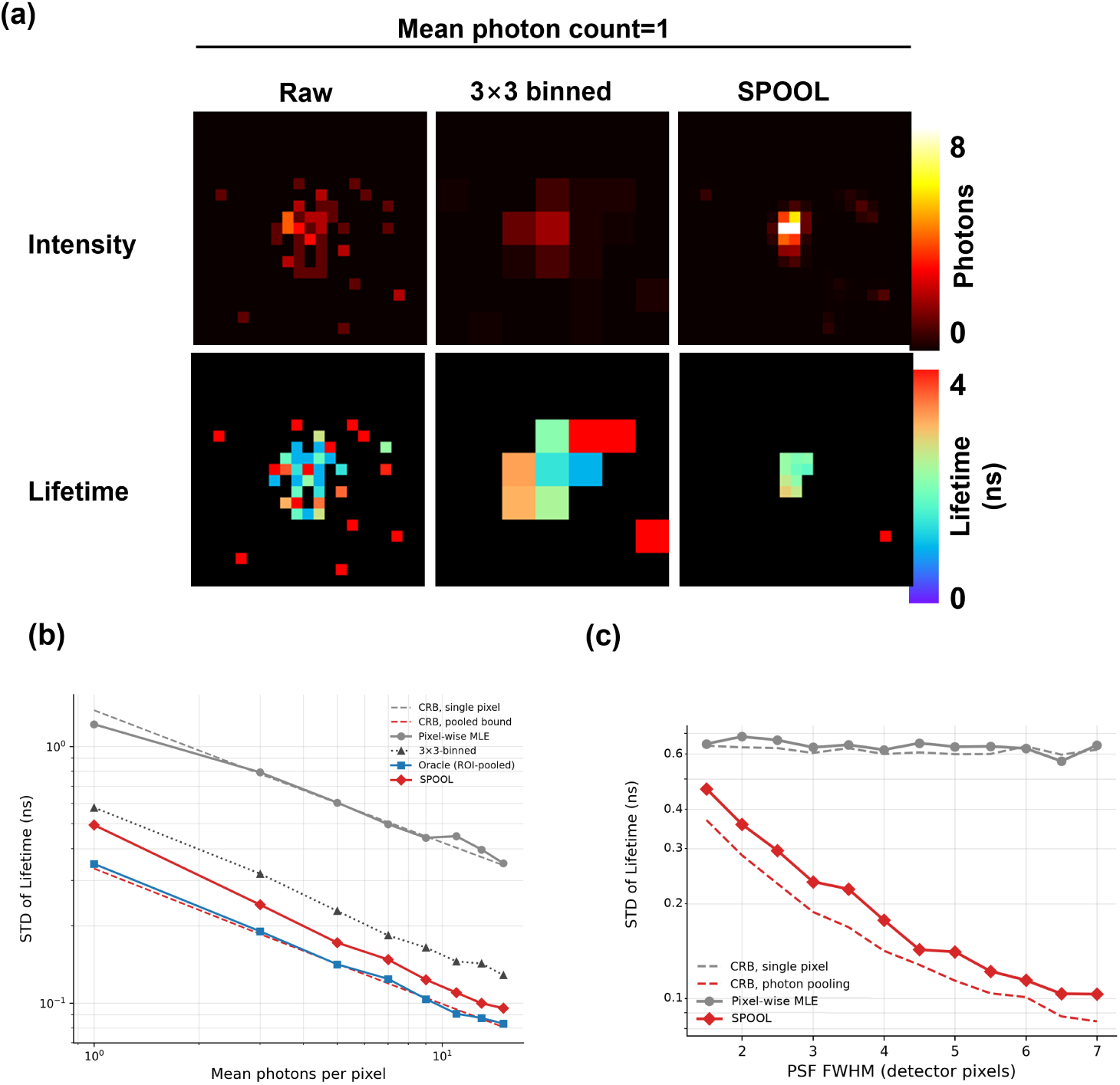
Joint inference approaches the pooled information limit without known source support. **a**, Raw measurement, 3 *×* 3 spatial binning and SPOOL for a single simulated emitter at a mean of one photon per foreground pixel. Top, intensity; bottom, recovered lifetime. **b**, Lifetime standard deviation across 400 independent Poisson realizations, for converged pixel-wise MLE, 3 *×* 3binned MLE, an oracle ROI-pooled MLE given the true emitter position and support, and SPOOL. Dashed curves denote the single-pixel and pooled Cramér–Rao bounds. **c**, Lifetime standard deviation versus PSF width at a fixed 200-nm PSF and five detected photons per foreground pixel, from 1.5 to 7 pixels per full width at half maximum. Dashed curves, the single-pixel and pooled Cramér–Rao bounds.

Figure 2b compares four estimators with the single-pixel and pooled Cramér–Rao bounds calculated using the same temporal response, background and acquisition window. The converged pixel-wise maximum-likelihood estimator followed the single-pixel bound across the photon sweep (median ratio, 1.01), showing that it efficiently used the photons available at an individual pixel. An oracle estimator given the emitter position and support, which sums the decay histograms over the PSF footprint and fits a single continuous lifetime, followed the pooled bound (median ratio, 1.02), confirming that the information distributed across the PSF was attainable when source assignment was known. A 3 × 3-binned estimator recovered a 2.7× gain, but at threefold coarser spatial sampling.

Without prior knowledge of the emitter position or support, SPOOL remained within approximately 20% of the pooled bound in standard deviation across the examined few-photon regime (median ratio, 1.18), that is, within 20% of the oracle that was given the source location. At photon levels of at least five photons per pixel, this corresponded to a 3.4–4.1 improvement over converged pixel-wise estimation, approaching the 4.3× value predicted from the optics alone. The reconstruction therefore recovered most of the information made available by the PSF while retaining a spatially resolved output.

The scale of the attainable gain is set by the optical sampling rather than selected as an algorithmic hyperparameter. Repeating the single-emitter analysis from 1.5 to 7 pixels per full width at half maximum at a fixed 200-nm PSF and five detected photons per foreground pixel demonstrates that the pixel-wise dispersion was independent of sampling to within 4%. This is because the central pixel receives the same photon count in every condition, whereas the joint estimate improved as *σ^−^*^1.00^ (Fig. 2c). The measured gain was accordingly proportional to the PSF width, with a fitted slope of 0.91 per pixel-per-FWHM (*R*^2^ = 0.98) against the 1.06 implied by Eq. (10). The residual difference is accounted for by how far each estimator sits from its own bound rather than by the prediction: their ratio, 1.04 for the pixel-wise estimator against its single-pixel bound and 1.21 for the joint estimator against the pooled bound, reproduces the measured slope. Relative to pixel-wise estimation, the lifetime standard deviation was reduced by 49% at two pixels per PSF full width at half maximum and by 84% at seven pixels per full width at half maximum. Below two pixels per full width at half maximum, the gain falls to 1.4×: a single pixel there already collects most of an emitter’s photons, and little delocalized information remains to be recovered.

Finer sampling does not create additional photons or increase the full-field Fisher information; it decreases the fraction captured by any one detector pixel and therefore enlarges the penalty incurred by pixel-wise estimation. Holding the mean count per foreground pixel fixed, as here, lets the total number of photons collected from the emitter grow as the sampling is refined, so the joint estimate improves while the pixelwise estimate does not. Holding the total photon budget integrated over the PSF fixed instead would leave the joint estimate unchanged while degrading the pixel-wise estimate. The ratio between the two, which is the quantity Eq. (10) predicts, is the same under either convention.

### 2.2 Joint inference remains accurate in multi-emitter scenes

The isolated-emitter analysis defines the information available when a source can be treated independently. We next tested whether this advantage persists in multi-emitter scenes, using a homogeneous phantom in which all emitters had a lifetime of 2 ns and a heterogeneous phantom containing emitters with lifetimes of 2 and 4 ns.

Representative reconstructions at 5.3 photons per foreground pixel are shown in Figs. 3a and 3c. Pixel-wise maximum-likelihood estimation (MLE), noise-corrected principal component analysis (NC-PCA) [34] and phasor analysis produced substantial variation within the diffraction-limited bead footprints. The few-photon fluorescence lifetime imaging (FPFLI) [4] network, retrained on data from this forward model, generated a more spatially coherent detector-space estimate but did not invert the acquisition PSF.

**Fig. 3.**
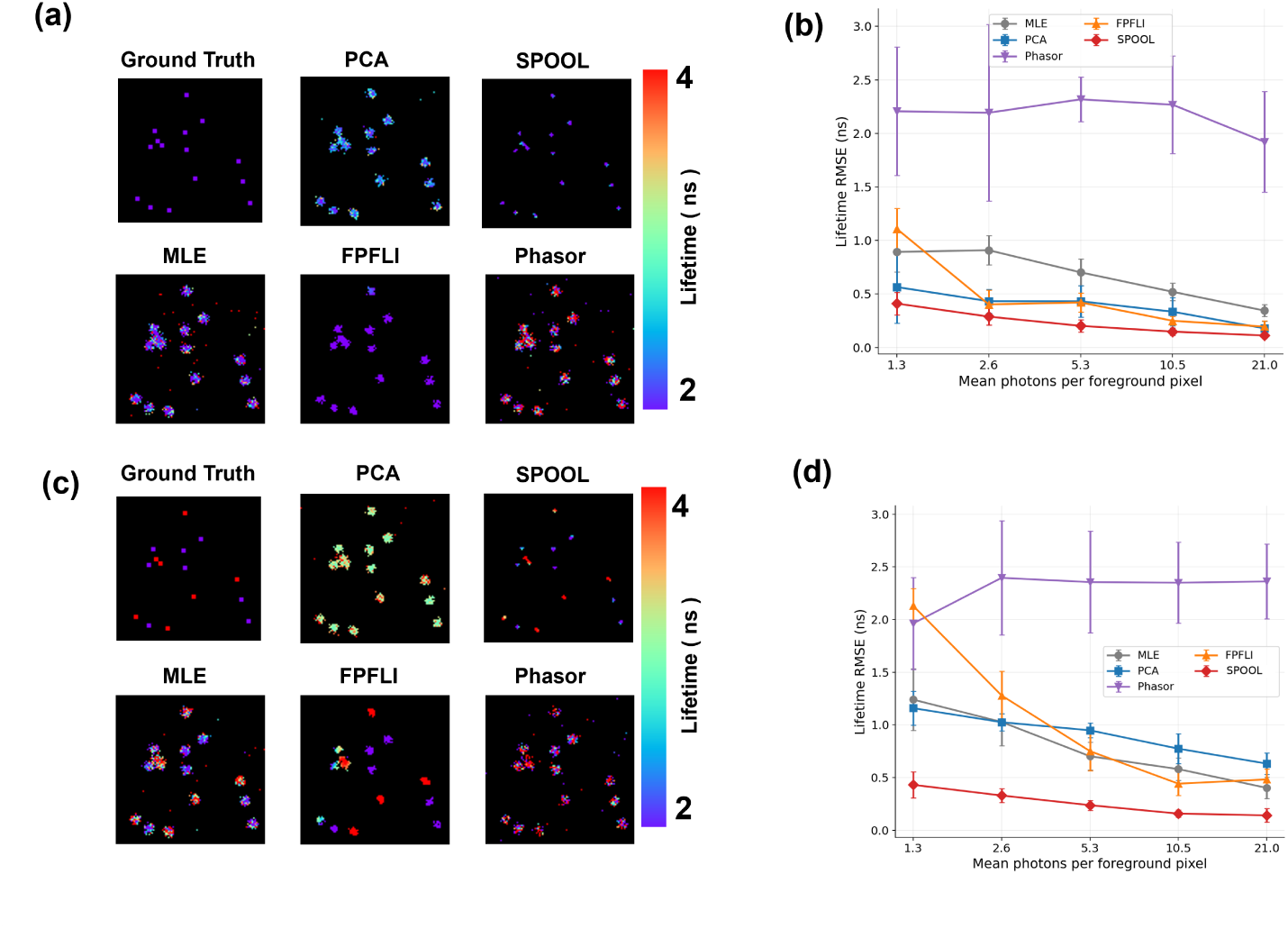
Simulation benchmark in bead phantoms. **a**, Representative lifetime maps for the homogeneous phantom (*τ* = 2 ns) at 5.3 detected photons per foreground pixel. In the ground-truth panel, each emitter occupies a single source pixel; for visibility only, emitters are displayed as 3 *×* 3-pixel markers centered on their true positions. This dilation is purely for display and enters neither the simulation nor any reconstruction or evaluation. **b**, Emitter-center lifetime RMSE for the homogeneous phantom as a function of the mean number of detected photons per foreground pixel. **c,d**, As **a,b** for the heterogeneous phantom (*τ* = 2 and 4 ns), with ground truth displayed as in **a**. FPFLI was retrained on data from this forward model. Curves show mean *±* s.d. across eight independent trials.

Quantitatively, SPOOL achieved the lowest lifetime root-mean-square error (RMSE) at every photon budget in both scenes (Figs. 3b,d). Relative to pixel-wise MLE it reduced the RMSE by 2.2–3.0-fold in the homogeneous scene and 2.0–3.1-fold in the heterogeneous scene. Against whichever baseline performed best at each budget the margin was 1.1–1.9-fold in the homogeneous scene and 2.0–3.1-fold in the heterogeneous scene: the separation is largest where the estimator must resolve contributions from emitters with different lifetimes within overlapping PSF footprints, which is the situation the joint likelihood is designed for. In the homogeneous scene at the highest budget examined the advantage over NC-PCA (0.169 versus 0.179 ns) lies within the trial-to-trial spread and should not be read as a difference.

FPFLI, like the 3 × 3-binned control of Fig. 2, increases the local photon budget through spatial pooling: its local lifetime estimator operates on 8 × 8 pooled blocks and recovers resolution afterwards by intensity-guided interpolation rather than by inverting the PSF. NC-PCA instead exploits low-rank structure across the temporal dimension while retaining the native spatial grid. Both close much of the gap at high photon budgets in the homogeneous scene, where local pooling or low-rank denoising is favorable, and neither does so in the heterogeneous scene, where overlapping lifetime species must be separated.

We next tested whether the improvement could be reproduced by performing contrast unmixing and spatial deconvolution sequentially. In the homogeneous phantom, three controls first unmixed each detector pixel using the same lifetime dictionary and then deconvolved the recovered amplitude maps using Wiener, total-variation or Richardson–Lucy reconstruction (Supplementary Fig. S1). Across the photon sweep, joint inference achieved a 2.0–2.5-fold lower RMSE than the best sequential control.

The Richardson–Lucy control used the same PSF, multiplicative update and iteration count, although its initial unmixing step remained least-squares based and therefore did not isolate operation order alone. Pixel-wise unmixing compresses the raw spatiotemporal photon distribution into independent, noisy amplitude estimates before any spatial information is used; joint inference, as in SPOOL, instead lets spatial assignment and contrast decomposition constrain one another within the Poisson likelihood.

Raw per-pixel phasor analysis produced the largest errors and yielded non-physical estimates for 18% of emitter centers in the homogeneous scene and 38% in the heterogeneous scene at the lowest photon budget. In the homogeneous scene these failures fell to 2% by the highest budget, whereas in the heterogeneous scene they persisted at 22–40% throughout, reflecting the difficulty of assigning a single phasor to a pixel containing two lifetimes. This result describes the deliberately unbinned phasor estimator used here; spatial pooling or filtering would improve its performance, but at the cost of spatial resolution.

### 2.3 Experimental photon efficiency on single-dye bead and microtubule samples

We next investigated whether the photon-efficiency advantage established in simulation survives under real experimental conditions. We begin with two single-dye samples, in which only one fluorophore is present: fluorescent bead standards, which present isolated, compact sources, and immunolabeled microtubules, which present an extended filamentous network. The two samples were acquired at pixel pitches of 30 and 39 nm, corresponding to 6.7 and 5.1 detector pixels per PSF full width at half maximum, and are analyzed at their native sampling.

Figures 4a and 4c show recovered lifetime maps at three thinned photon budgets (1, 5, and 15 mean photons per foreground pixel). At 1 photon per pixel, pixel-wise MLE produces a lifetime map dominated by pixel-to-pixel scatter, in which the individual beads and the filament network are visible only as noisy silhouettes. SPOOL recovers, from the same photons, compact beads and a continuous, legible filament network – structure that the pixel-wise estimate does not resolve at this photon budget.

**Fig. 4.**
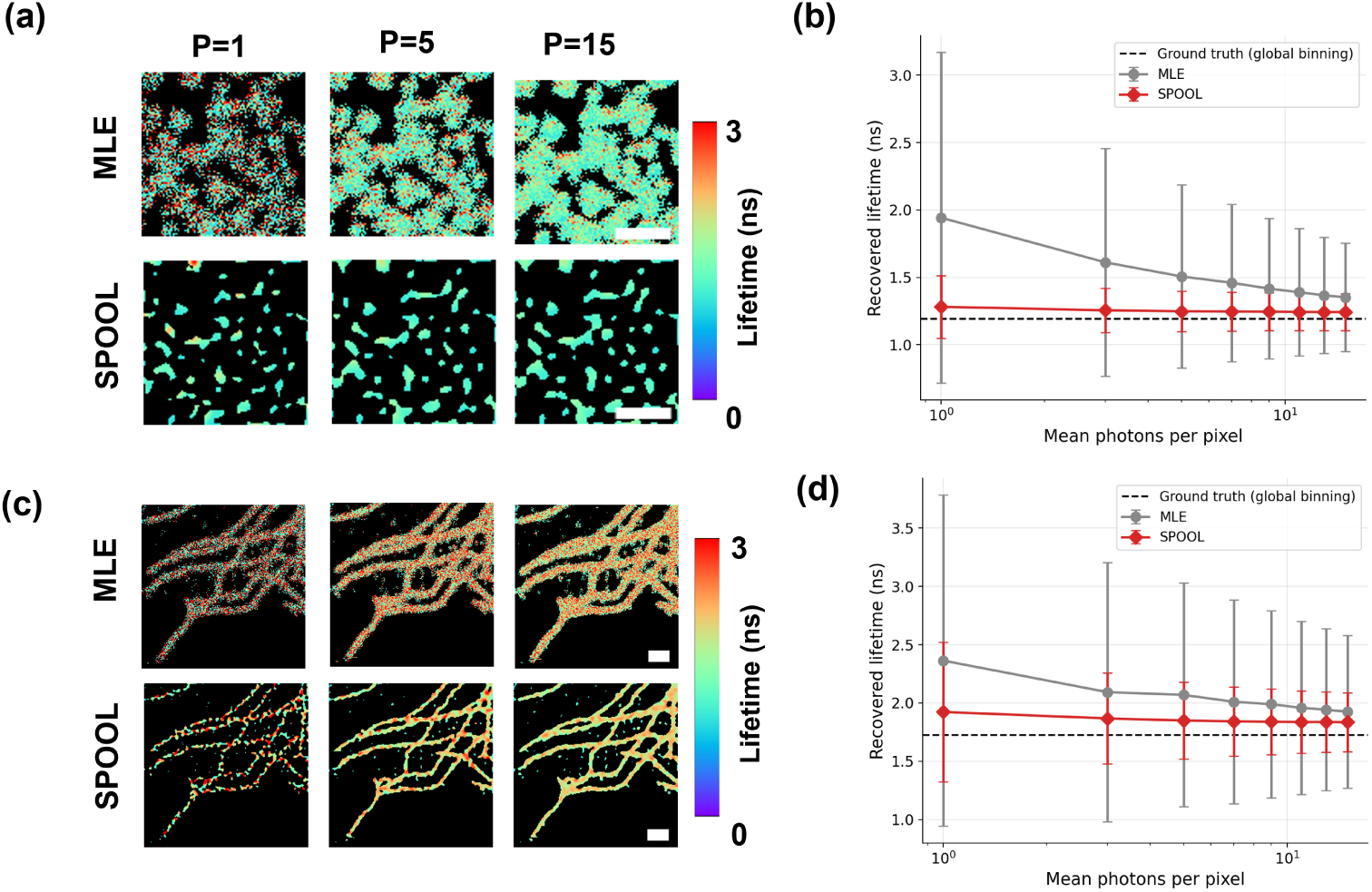
Photon efficiency in single-dye experimental samples. **a**, Lifetime maps of fluorescent beads using pixel-wise Poisson MLE (top) and SPOOL (bottom). **b**, Recovered lifetime for fluorescent beads as a function of the mean number of detected photons per foreground pixel. Points show the mean recovered lifetime and error bars the s.d. across six independent thinnings at each photon level; the dashed line denotes the globally pooled full-photon estimate. **c**, Lifetime maps of Oregon Green-labelled microtubules using pixel-wise Poisson MLE (top) and SPOOL (bottom). **d**, Recovered lifetime for microtubules, plotted as in **b**. Scale bars, 1 µm.

Figures 4b and 4d quantify this across a sweep from 1 to 15 mean photons per pixel. For the bead sample, SPOOL’s standard deviation was 0.194 ns at 1 photon per pixel, against 1.165 ns for pixel-wise MLE, a 6.0-fold improvement. The microtubule sample follows the same pattern with a somewhat smaller margin, as expected for an extended structure in which neighboring pixels share less independent information than isolated beads do: 0.442 versus 1.354 ns at 1 photon per pixel, a 3.1-fold improvement.

### 2.4 Experimental photon efficiency on a dual-labeled, two-species cell sample

We next tested a more demanding, biologically realistic case in which two distinguishable lifetime species (mitochondria labelled with Alexa Fluor 488 and microtubules with Alexa Fluor 555) coexist within the same field of view and are frequently blended within individual, PSF-blurred pixels.

Figure 5a shows each method’s reconstruction from the full, unthinned acquisition, which serves as that method’s own high-photon reference. Low-photon measurements were generated from the same raw photon-count cube by binomial photon thinning [35]: each detected photon was independently retained with probability *p* = *n/n*_full_, where *n* is the target mean photon count per foreground pixel. Because binomial thinning of a Poisson process preserves Poisson statistics, this procedure reproduces the statistics expected from a shorter acquisition of the same specimen. Further details on the photon-thinning procedure are provided in Supplementary Section S6. Figure 5b reports the reference-relative RMSE across a 1–15 photon-per-pixel sweep via the photon-thinning process. Figures 5c and 5d show the recovered maps and per-pixel absolute reference-relative error at 1 and 5 photons per pixel. At 1 photon per pixel, pixel-wise MLE yields a map in which pixel-level noise obscures essentially all spatial structure; SPOOL instead recovers a map in which the punctate mitochondrial network remains legible against the surrounding microtubule-dominated background.

**Fig. 5.**
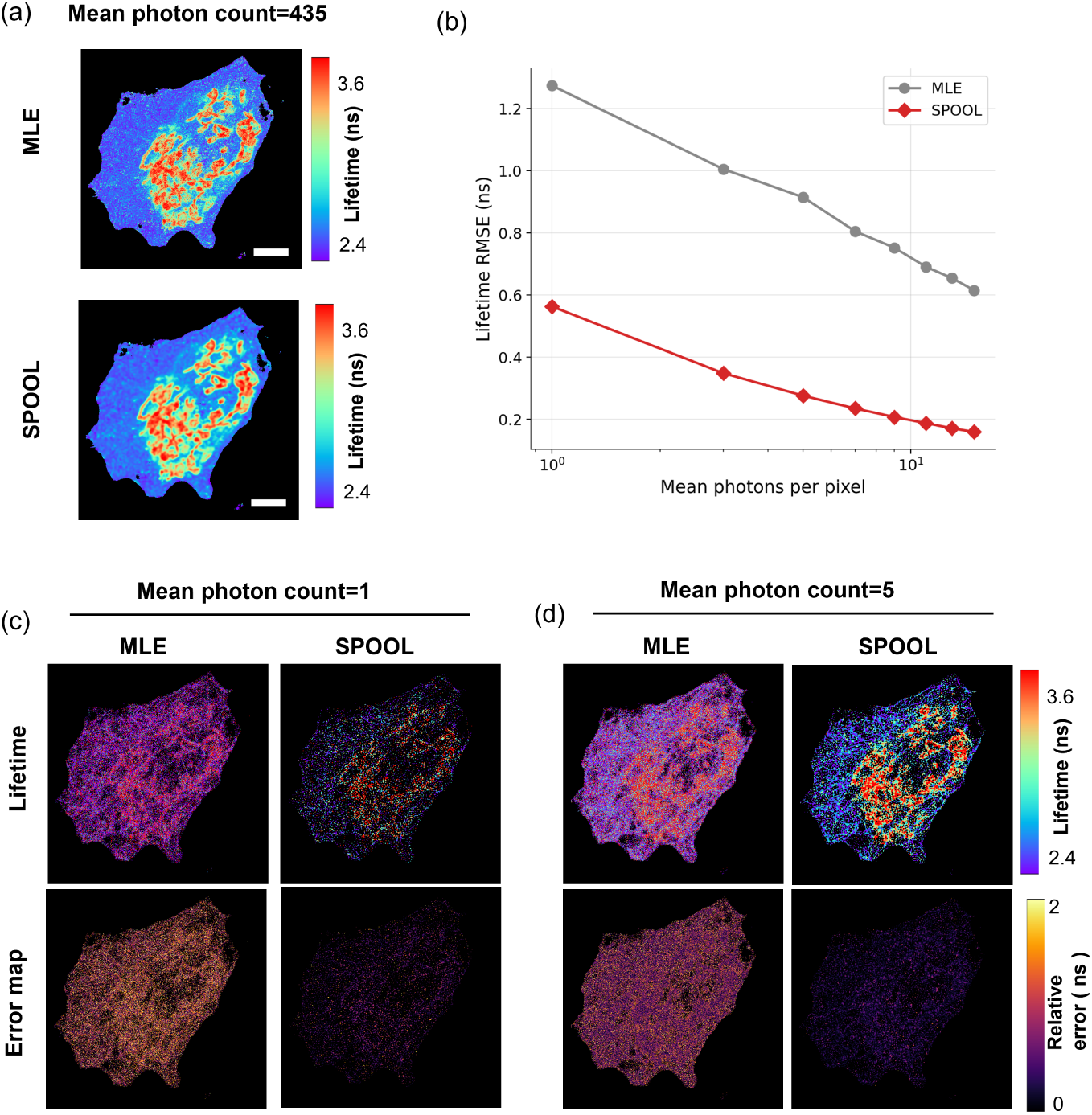
Photon efficiency on a dual-labeled cell sample. (mitochondria: Alexa Fluor 488; microtubules: Alexa Fluor 555). **a**, Full-photon lifetime maps from pixel-wise Poisson MLE (top) and SPOOL (bottom). **b**, Reference-relative lifetime RMSE for pixel-wise MLE and SPOOL as a function of the mean number of detected photons per foreground pixel, computed by pooling errors across six independent photon thinnings and all pixels in each method’s recovered support. **c,d**, Lifetime maps (top) and absolute reference-relative error (bottom) at 1 and 5 photons per foreground pixel, respectively. Scale bars, 5 µm.

Quantitatively, the reference-relative lifetime RMSE was 1.192 ns for pixel-wise MLE versus 0.447 ns for SPOOL at 1 photon per pixel, a 2.7-fold reduction, widening to 3.3-fold at 3 photons, 3.8-fold at 5, and 4.3-fold at 15 photons per pixel (0.549 versus 0.127 ns). Equation (10) predicts 3.6× from this sample’s optics alone (*σ*_PSF_ = 1.41 px at the 60.4 nm pixel size), and the measured range straddles it.

The two methods differ in their *mean signed reference error* and not only in their dispersion, which is why RMSE rather than the standard deviation alone is the appropriate summary here (Methods). At 1 photon per pixel the pixel-wise estimate sits 0.279 ns below its own full-photon answer and is still 0.030 ns below it at 15, whereas SPOOL is within 0.024 ns at 1 photon and 0.002 ns at 15. The pixel-wise estimator is therefore not merely noisier at low photon counts; it approaches its own high-photon limit from a systematic offset that decays slowly with photon number.

### 2.5 The optics-defined gain transfers to hyperspectral imaging

The Fisher-information analysis does not depend on excited-state decay. It requires only a calibrated spatial forward model and a contrast-axis dictionary. We therefore replaced the temporal decay profiles with *K* = 11 normalized Gaussian emission bands spanning 532–632 nm (Supplementary Table S2). No fluorophore-specific spectrum or identity was supplied. The PSF, Poisson likelihood, multiplicative update, damping and iteration count were unchanged, yielding a source-space mean-wavelength map in place of a lifetime map.

The hyperspectral reconstruction reproduced the few-photon behaviour observed in FLIM (Fig. 6). At one photon per foreground pixel, the pixel-wise estimate was dominated by spectral scatter, whereas the joint reconstruction recovered the filamentous and punctate structures and began to distinguish their spectral populations (Fig. 6c). At five photons per pixel, the two populations were clearly separated while the pixel-wise map remained noisy (Fig. 6d). Relative to each method’s full-photon reconstruction, the wavelength RMSE decreased from 26.36 to 11.13 nm at one photon per pixel and from 10.14 to 2.51 nm at 15 photons per pixel, corresponding to 2.4and 4.0-fold improvements, respectively. Because each method is evaluated over its own recovered support, the two masks contain different pixel counts by construction: 1.5 10^5^ pixels for pixel-wise MLE versus 6.3 10^4^ for SPOOL at one photon per pixel, converging toward 2.0 10^5^ versus 1.3 10^5^ at 15.

**Fig. 6.**
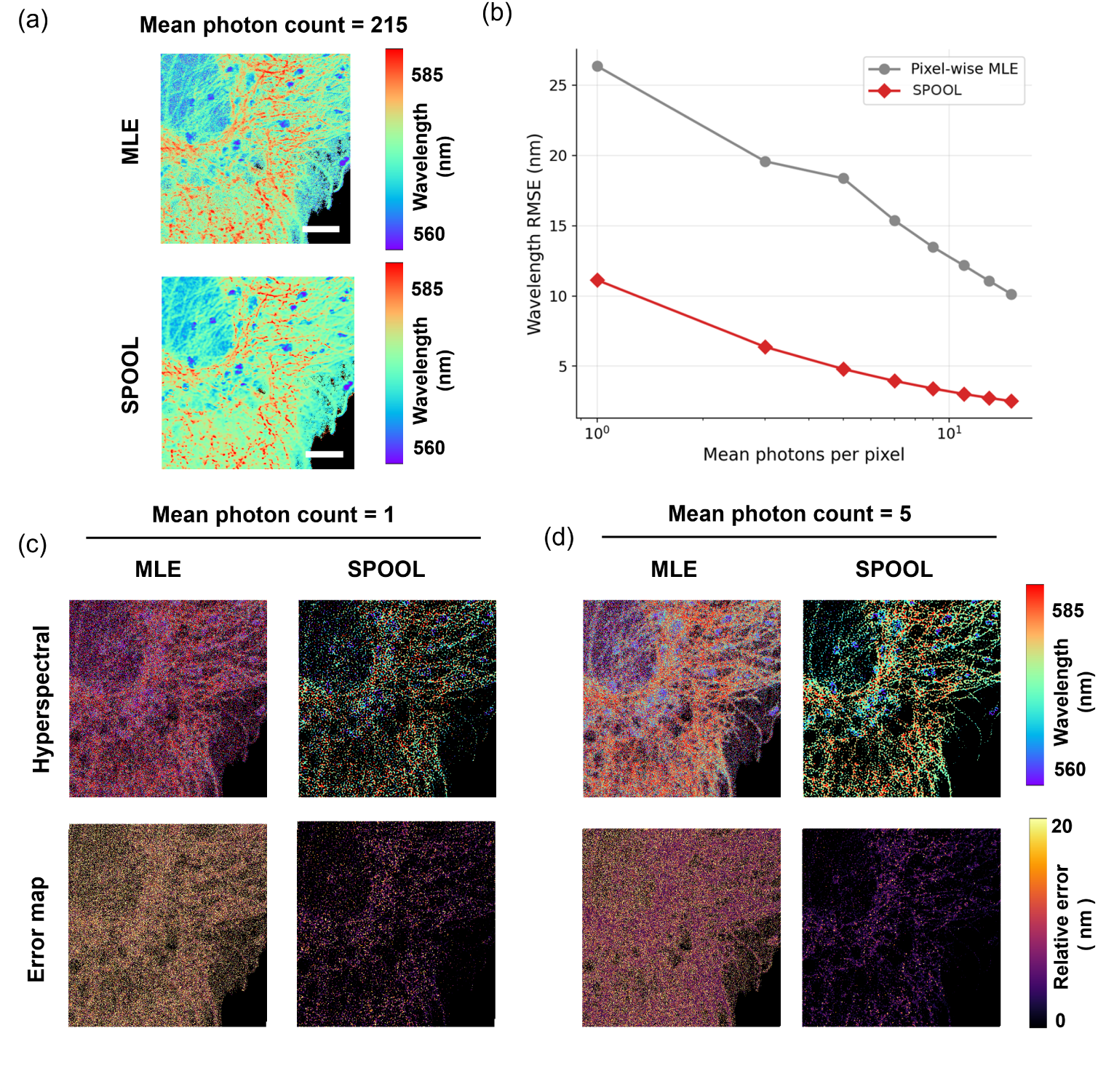
Few-photon hyperspectral imaging. **a**, Full-photon (215 per pixel) mean-wavelength map. **b**, Reference-relative wavelength RMSE as a function of the mean number of detected photons per foreground pixel. **c,d**, Mean-wavelength maps (top) and absolute errors (bottom) at 1 and 5 photons per pixel, respectively. Scale bars, 5 µm.

This sample was acquired at the same 50 nm sampling as the simulations, giving *σ*_PSF_ = 1.70 pixels and an optics-derived prediction of 4.3×. The measured gain rose from 2.4× at one photon per pixel to 4.0× at 15, approaching this prediction from below as the photon budget increased — the same behaviour observed for the dual-labeled FLIM sample — despite the change from temporal to spectral contrast, supporting the proposed mechanism across a second contrast axis. The photon dependence also differed between the estimators. Across the measured range, the reference-relative RMSE of the joint reconstruction followed an empirical scaling of *n^−^*^0.55^, close to the *n^−^*^1*/*2^ dependence expected for shot-noise-limited estimation, whereas the pixel-wise reconstruction exhibited a shallower dependence of *n^−^*^0.35^. The shallower exponent is consistent with the pixel-wise estimator remaining in a low-count, non-asymptotic regime, while spatial pooling gives the joint estimator a larger effective photon budget per estimate and can bring it closer to the asymptotic regime at lower counts. These exponents describe the observed error scaling over the finite photon range examined, not a change in the fundamental shot-noise limit.

## 3 Discussion

Photon starvation in quantitative fluorescence microscopy is usually framed as an acquisition problem: too few photons reach each pixel to support a reliable estimate. Our results show that this can be cast as an inference problem. The microscope optics distribute photons from one source across several detector pixels, yet conventional analysis asks each pixel to estimate molecular contrast independently. By restoring this spatial relationship within the likelihood, SPOOL retains quantitative information under photon budgets at which pixel-wise inference becomes unstable. For an isolated emitter sampled at four pixels per PSF full width at half maximum, the optics-derived theory predicted a 4.3-fold precision gain over single-pixel estimation, and the reconstruction recovered most of this information without being given the emitter position or support. The advantage persisted when sources overlapped and carried different lifetimes, for which pooling regions cannot be defined in advance. At one detected photon per foreground pixel, the method reduced lifetime dispersion sixfold in fluorescent-bead experiments and reduced the lifetime root-mean-square error relative to a high-photon reference from 1.19 to 0.45 ns in dual-labeled cells. The central result is therefore not simply that pooling photons improves precision, but that much of this information can be recovered by incorporating the spatial constraints imposed by the imaging PSF into the likelihood, without requiring prior knowledge of emitter positions or predefined pooling regions, while retaining spatially resolved quantitative contrast.

The work builds on a broader effort to incorporate image-formation physics into computational microscopy. Previous methods have encoded molecular dynamics in spatial patterns or recovered information distributed across detector coordinates by explicitly modelling multidimensional image formation [24–26], incorporated physical forward models into learned reconstruction [20, 21], and improved spectral decomposition at low signal by accounting for noise and spectral structure [1, 3]. Here, the complementary objective is to use spatial information imposed by the PSF to improve estimation along a temporal or spectral contrast axis, solving image formation and contrast decomposition jointly so that photons detected at different pixels constrain the same source-space components. The forward model therefore does not serve only as a prior or training constraint; it directly defines the estimator and predicts the magnitude of the attainable gain.

The information-bound analysis clarifies the origin and the scope of this gain. When the position and support of an isolated emitter are known, summing its photons is sufficient to approach the pooled Cramér–Rao bound, and the oracle estimator confirms that this limit is physically attainable. The oracle does not, however, solve the general imaging problem: it requires source locations and extents in advance, returns one parameter per predefined region rather than a spatial map, and cannot assign photons unambiguously when sources overlap. SPOOL instead reconstructs a complete source-space field with unknown source locations, a finite contrast dictionary and nonnegativity constraints, and its standard deviation remained within approximately 20% of the pooled bound across the examined few-photon regime. The converged pixel-wise estimator, by contrast, closely followed its own single-pixel bound and was therefore efficient for the problem it was asked to solve; its limitation was not poor optimization but incomplete use of the measurement. The gain arises because the estimation problem has been reformulated, not because an inefficient algorithm has been replaced by a better optimizer. This perspective complements recent efforts to evaluate computational microscopes according to how closely they approach the precision permitted by their measurements, rather than by image quality alone [36].

Joint inference is also not equivalent to deconvolving an already estimated parameter map. In the closest sequential control, the same PSF, Poisson multiplicative update and iteration count were applied after pixel-wise unmixing, yet joint inference remained 2.0–2.5-fold more accurate across the photon sweep. Once each detector pixel has independently converted its sparse decay or spectrum into component estimates, the relationship among photons distributed across neighboring pixels has been compressed into separate, heteroscedastic estimates; subsequent deconvolution can sharpen these maps but cannot restore that relationship. In the joint likelihood, spatial reassignment and contrast decomposition constrain one another before either decision is made.

The hyperspectral experiment shows that the same mechanism extends beyond lifetime contrast. The temporal-decay dictionary was replaced by generic Gaussian spectral bands without changing the spatial model, likelihood or optimization and without providing fluorophore identities or measured emission spectra. At one photon per foreground pixel the wavelength root-mean-square error relative to the high-photon reconstruction decreased from 26.36 to 11.13 nm, and by five photons per pixel the error fell below the separation between the recovered spectral populations. The gain therefore does not require lifetime-specific photophysics: the contrast axis determines the absolute information carried by each photon, whereas the relative advantage obtained by recovering spatially distributed photons is governed by the optical sampling. The framework may extend to other photon-counting measurements represented by a calibrated spatial operator and a non-negative linear contrast basis, including time-gated and polarization-resolved imaging, whereas nonlinear image formation, coherence or substantially non-Poisson statistics will require different forward models. Three factors bound the scope of these results. First, the method assumes a known, spatially invariant PSF, set here from the objective, representative wavelength and pixel pitch rather than fitted to each dataset. Reconstructions with deliberately misspecified optics retained at least 86% of the measured gain across a ±20% error in the assumed PSF width, approximately twice the variation expected across the relevant emission band, and never fell to the pixel-wise baseline (see Supplementary Information); experimentally measured, wavelength-dependent and spatially varying PSFs are natural extensions. Second, the predicted gain depends on the PSF width in detector pixels and is consequently a property of the acquisition sampling as well as of the estimator. The datasets examined here span 3.3 to 6.7 pixels per full width at half maximum, with corresponding predicted gains of 3.6–7.1-fold, and at coarser sampling the gain approaches unity because a single detector pixel then collects most of an emitter’s photons. It is by contrast relatively insensitive to background: reducing the foreground signal-to-background ratio from 50 to 5 changes it by 7%, and by 2% once expressed as a fraction of the gain attainable at that ratio (see Supplementary Information). These analyses assume a spatially homogeneous background; spatially heterogeneous backgrounds were not modeled here and may require explicit background estimation or an extended forward model. Extended or densely overlapping structures need not attain the isolated-emitter value, as reflected by the smaller improvements measured for microtubules. Third, the iteration count acts as an implicit regularization parameter because the multiplicative reconstruction is semi-convergent. The RMSE-minimizing stopping point varies with photon level, so no single iteration count is optimal at every photon budget. Rather than tuning the reconstruction separately at each photon level, we fixed the iteration count at 50, which lies within the broad minimum of the RMSE averaged over 1–15 photons per pixel and remains near-optimal when the assumed PSF width is perturbed by ±10% (Supplementary Section S7). This fixed operating point avoids photon-level-specific tuning while retaining low error across the regime examined here. Additional PSF- and background-robustness analyses are provided in Supplementary Section S8, and runtime and memory benchmarks are provided in Supplementary Section S9. Automated stopping criteria or explicit regularization may further improve performance when absolute parameter accuracy is the primary objective.

The reconstruction is computationally practical: at the fixed 50-iteration setting with *K* = 11, all experimental datasets reconstructed in under 2 s using double-precision computation and under 0.35 s using single precision on a single NVIDIA GeForce RTX 3060 Ti, with peak GPU memory usage below 1.9 GiB. The same training-free multiplicative update was used across all datasets without retraining for a particular specimen or contrast modality, making the framework compatible with downstream processing of existing photon-counting measurements. Taken together, these results show that the precision of a quantitative optical measurement is determined not only by how many photons are detected, but also by whether the estimator preserves the relationships that the instrument has imposed between them. The PSF is usually treated as a loss of spatial resolution. Here, it also serves as a map linking photons that carry information about the same underlying source. By incorporating that map before contrast estimation, SPOOL recovers quantitative lifetime and spectral information without known source assignments and under photon budgets at which conventional pixel-wise analysis becomes unstable. Some information that appears to be absent has instead been spatially redistributed by the optics. Recovering that information provides a physics-defined route to few-photon quantitative imaging.

## Data availability

The experimental photon-count data supporting this study are available from the corresponding author on reasonable request.

## Code availability

SPOOL is publicly available as a Fiji plugin through the SPOOL update site. The source code and analysis scripts used to produce the results reported in this study are publicly available at https://github.com/Wonsang7/SPOOL under the MIT License. The repository includes the simulation benchmarks and analysis pipelines for the FLIM and hyperspectral imaging experiments presented in this study.

## Acknowledgements.

I.C.H. is supported by the National Institute of Biomedical Imaging and Bioengineering under award number 1K25EB032864 and by a Physician/Scientist Development Award from Massachusetts General Hospital. W.H. and C.L.E. are supported by the Ludwig Cancer Center at Harvard.

## Author contributions

W.H. conceived the study, developed the method, implemented the simulations and reconstruction software, analyzed the experimental data, and wrote the original draft of the manuscript. I.C.H. acquired the experimental data and contributed to reviewing and editing the manuscript. C.L.E. provided resources and supervision and contributed to reviewing and editing the manuscript. All authors read and approved the final manuscript.

