## Supplementary Information for "Spatially pooling photon information enables photon-efficient quantitative imaging"

Every value reported below was read out of the code that produced the figures. Equation and section numbers prefixed with S refer to this document; unprefixed references are to the main text.

#### S1 Reconstruction implementation

Every SPOOL reconstruction in this work – FLIM and hyperspectral, simulated and experimental – uses the same solver. The only inputs that differ between datasets are the PSF width, the dictionary, the background level, and the data themselves.

***Initialization.***

The amplitude maps are initialized uniformly at $\max\left(\sum_{\mathbf{r},t} Y(\mathbf{r},t)/(HWK),\, 10^{-4}\right)$ for a field of $H \times W$ pixels and $K$ components: a flat map carrying the measured total photon count, chosen so that the first iteration is not dominated by an arbitrary scale.

***Numerical safeguards.***

A floor $\epsilon = 10^{-9}$ is added to every denominator. The multiplicative update ratio $U_k/(C_k + \epsilon)$ is clipped to be nonnegative *before* the fractional damping exponent $\eta$ is applied. This clip is not cosmetic: floating-point round-off in the spatial convolution can occasionally produce a formally negative base for a fractional power, and the resulting not-a-number value is then absorbed at zero by the non-negativity floor and, given the purely multiplicative nature of the update, collapses the entire reconstruction irrecoverably. The amplitude estimate is floored at $10^{-8}$ rather than exactly zero at each iteration, so that a pixel driven to zero can recover.

***Spatial convolution and its adjoint.***

The convolutions $h * A_k$ and $h^{\mathrm{T}} * (\cdot)$ are evaluated by FFT, zero-padded to the full linear-convolution size and cropped back, i.e. with a zero boundary condition. A plain FFT product would implement a *circular* convolution, wrapping the opposite edge of the field around, and would differ from the intended operator at the image borders. The adjoint uses the transform of the flipped kernel, $h(-\mathbf{r})$, rather than the conjugate transfer function, which avoids an off-by-one shift in the cropping window. We verified the implementation against a direct spatial convolution to a relative accuracy of $10^{-15}$, and verified the adjoint identity $\langle h * a,\, b\rangle = \langle a,\, h^{\mathrm{T}} * b\rangle$ to a relative accuracy of $10^{-10}$.

***Point spread function.***

The PSF is a two-dimensional Gaussian *integrated over each detector pixel* rather than sampled at pixel centers,

$$h(i,j) \;\propto\; \Big[\mathrm{erf}\big(\tfrac{i+1/2}{\sqrt{2}\sigma_{\mathrm{PSF}}}\big) - \mathrm{erf}\big(\tfrac{i-1/2}{\sqrt{2}\sigma_{\mathrm{PSF}}}\big)\Big]\Big[\mathrm{erf}\big(\tfrac{j+1/2}{\sqrt{2}\sigma_{\mathrm{PSF}}}\big) - \mathrm{erf}\big(\tfrac{j-1/2}{\sqrt{2}\sigma_{\mathrm{PSF}}}\big)\Big], \tag{S1}$$

normalized to unit sum and truncated at $\pm 4\sigma_{\mathrm{PSF}}$. Pixel integration matters because $h_0$, the fraction of an emitter's photons falling in its central pixel, is the quantity that sets the predicted gain: point sampling misestimates it by 0.5% at $\sigma_{\mathrm{PSF}} = 2.8$ px and by 4% at $\sigma_{\mathrm{PSF}} = 1.0$ px.

***Background.***

$B(\mathbf{r}, t)$ is taken to be spatially and temporally uniform and estimated from a $30 \times 30$ pixel corner region of the field containing no labeled structure,

$$B \;=\; \frac{1}{900\,T} \sum_{\mathbf{r}\in\mathrm{corner}} \sum_{t=1}^{T} Y(\mathbf{r}, t), \tag{S2}$$

the mean counts per pixel per bin in that region. The same construction is used over the spectral axis for the hyperspectral data. Under photon thinning at retention fraction $p$ the background is thinned with the data, $B \to pB$, since the detector thins background counts too. Omitting this scaling would leave a background that is too bright relative to the thinned signal at low photon levels and would inflate the apparent low-photon error of every method. The two simulations set the background differently, by design. The single-emitter study uses a fixed $B = 10^{-4}$ counts per pixel per time bin — 0.026 photons per pixel over the 256-bin window, about 2% of the signal at one detected photon per foreground pixel — because it tests whether the information limit of Eq. (10), which is derived in the background-free limit, is attainable in practice; a larger background would confound the efficiency of the estimator with the background level. The multi-emitter benchmark instead scales $B$ proportionally with the foreground signal at each photon level, maintaining a mean foreground-to-background ratio of 50. This matches photon thinning, which reduces signal and background counts by the same fraction and therefore preserves their relative ratio. Both are conservative against the measurement: the 114 time bins preceding the excitation pulse in the microtubule acquisition contain no counts at all across the entire field, bounding its background below $2.4 \times 10^{-6}$ counts per pixel per bin. Sec. S8 sweeps this ratio from 50 down to 5 and reports the consequences for both settings.

***No post hoc filtering.***

No spatial binning, median filtering, smoothing, or denoising was applied to the reconstructed parameter maps. Spatial coupling enters only through the PSF term in the raw photon-count forward model. Source-space maps were computed directly from the recovered amplitude maps; detector-space maps were obtained only by explicit forward projection with the same PSF.

### S2 Dictionary and IRF construction

***Instrument response.***

The IRF was not measured separately but estimated from the data, by fitting a Gaussian-convolved single-exponential decay (a reconvolution model) to the spatially-integrated decay curve. The fit returns a temporal offset $t_0$ and a Gaussian width $\sigma_{\mathrm{IRF}}$ (Table S1). The same $(t_0, \sigma_{\mathrm{IRF}})$ then enter every basis in the decay dictionary, so the temporal offset is handled inside the dictionary rather than by shifting the data. The fitted values were used only as shared calibration parameters of the dictionary and were not adjusted to optimize reconstruction performance. Because a reconvolution fit trades the IRF width against the recovered lifetime, this estimate is less well constrained for the dual-labeled sample, in which more than one lifetime contributes to the integrated decay. An independently measured IRF could further reduce this source of uncertainty.

***Decay dictionary.***

Each basis is the analytic convolution of a Gaussian IRF with a single exponential,

$$D_k(t) \;\propto\; \tfrac{1}{2}\lambda_k \exp\!\Big(\tfrac{1}{2}\lambda_k^2\sigma_{\mathrm{IRF}}^2 - \lambda_k(t-t_0)\Big)\,\mathrm{erfc}\Big(\frac{t_0 + \lambda_k\sigma_{\mathrm{IRF}}^2 - t}{\sqrt{2}\,\sigma_{\mathrm{IRF}}}\Big), \qquad \lambda_k = 1/\tau_k, \tag{S3}$$

clipped at zero and normalized to unit area. The unit-area normalization matters: without it the amplitude-weighted parameter map defined in the main text would not be physically meaningful, because bases with different $\tau_k$ would carry different total weight.

The dictionary is *lifetime-agnostic* throughout: $K = 11$ bases spanning 1.0–5.0 ns in 0.4 ns steps, chosen to cover the plausible range without fixing the reconstruction to the nominal lifetime of any particular dye. The same lifetime-agnostic dictionary is used for the experiments, the single-emitter study, and the multi-emitter benchmark so that no reconstruction method is provided with the ground-truth lifetimes. Scene generation uses the two true lifetime components only to define the ground truth; no reconstruction is given those components.

***Spectral dictionary.***

The hyperspectral reconstruction uses the direct analogue on the wavelength axis: $K = 11$ Gaussian emission bands with centers spaced 10 nm apart from 532 to 632 nm and a 40 nm FWHM each, normalized to unit area and constructed to tile the detection window. *No fluorophore spectrum is used.* The published emission spectra of the two dyes enter neither the model nor the fit, exactly as no dye lifetime enters the FLIM dictionary. For the parameter map defined in the main text, $\theta_k$ is set to the center wavelength $\lambda_k$ of the $k$-th band, so the recovered map is an amplitude-weighted mean emission wavelength.

One consequence should be stated: because the dictionary tiles the whole detection band, the recovered mean wavelength is a compressed contrast. Two dyes whose emission peaks differ by tens of nanometres can yield recovered means that differ by considerably less. This limitation is discussed in the hyperspectral Results section of the main text.

### S3 Single-emitter simulation and oracle estimator

A single point emitter was placed at the center of a $48 \times 48$ pixel field ($2.40 \times 2.40$ µm) with a 50 nm pixel size and a pixel-integrated Gaussian PSF of 200 nm FWHM ($\sigma_{\mathrm{PSF}} = 1.699$ px, four pixels per FWHM), with 256 time bins of 0.0977 ns, a Gaussian IRF of 150 ps full width at half maximum ($\sigma_{\mathrm{IRF}} = 0.064$ ns) at $t_0 = 0.50$ ns, within the 116–157 ps range measured on the experimental data, and a uniform dark-count level of $10^{-4}$ photons per pixel per time bin. This sampling matches the hyperspectral acquisition and corresponds to the confocal Nyquist rate $\lambda/8\mathrm{NA} \approx 50$ nm for these optics. The true lifetime was $\tau_0 = 2.0$ ns, placed *exactly midway* between the two nearest nodes of the reconstruction dictionary (1.8 and 2.2 ns; grid 1.0–5.0 ns in steps of 0.4 ns), the least favorable position for grid quantization; placing the truth on a grid node would have zeroed the quantization bias artificially.

At each photon level, 400 independent Poisson realizations were drawn and every estimator applied to each. Bias, standard deviation and root-mean-square error were computed *across realizations at the emitter pixel*, so that the measured variance is the replicate variance at a fixed parameter rather than a dispersion across pixels of differing brightness. Photon levels are quoted as the mean count per pixel over the emitter's foreground (the 37 pixels above 15% of the peak), the same rule used for the experimental data, so that the horizontal axis is directly comparable with the experimental figures. The ratio between the central pixel and this foreground mean is fixed by the PSF geometry at 2.3, so a mean of one photon per pixel places 2.3 photons in the central pixel and 43 photons across the full footprint.

Four estimators were compared.

***Pixel-wise Poisson MLE, converged.***

The pixel-wise form of the main-text multiplicative update with $h \to \delta$ and $\eta = 1$, run to convergence (1000 iterations; the change in $\hat{\tau}$ over the final 300 iterations was below $10^{-3}$ ns). Running this baseline to convergence rather than at the 50 iterations used for SPOOL is deliberate. Early stopping is a regularizer for an *ill-posed* deconvolution, and a pixel-wise Poisson fit is not ill-posed, so there is no basis for stopping it early. An under-iterated pixel-wise fit is a shrinkage estimator whose variance is artificially suppressed; using one as the baseline would both understate the reported gain and place the baseline, without justification, below its own Cramér–Rao bound. Each estimator is therefore run under the stopping rule appropriate to it.

$3 \times 3$-***binned pixel-wise MLE.***

What a practitioner actually does at these photon counts: bin the data spatially, then fit. Included because it is the obvious objection to the whole approach, and because it costs a factor of three in spatial resolution. The binning grid is offset so that the emitter falls at the *center* of its own bin. With an arbitrary grid phase the emitter lands on a bin edge and the central bin captures 30% rather than 39% of its photons, a 30% handicap that is an artifact of grid alignment rather than a property of binning.

***Oracle ROI-pooled MLE.***

Told the emitter's exact position, this estimator sums the decay histograms over a disc of radius $3\sigma_{\text{PSF}}$ around it – 81 pixels, capturing 98.6% of the emitter's photons – into a single histogram, and fits the two parameters $(N, \tau)$ by continuous Poisson maximum likelihood, with no dictionary, no deconvolution and no spatial regularization. The amplitude is profiled out by Newton iteration at each candidate lifetime rather than approximated by the background-subtracted total count, which would bias the oracle away from its own bound once the background is a few percent of the signal. This is the estimator that the pooled bound actually describes. It is included for two reasons. First, it establishes that the bound is attainable in this setting, which is what licenses using it as a benchmark at all. Second, it is an *oracle*: it must be told where the emitter is and where its support ends, it returns a single number rather than a map, and it cannot be run on an extended object. The gap between it and SPOOL is therefore the price of not being given that knowledge, and of returning a spatially resolved reconstruction instead.

***Proposed method.***

The full update with the PSF term, $\eta = 0.9$, 50 iterations, uniform initialization.

***Bounds.***

The dashed curves in the main-text single-emitter figure are Cramér–Rao bounds for the two-parameter model $(N, \tau)$, with a continuous lifetime and a known emitter position, evaluated numerically from the same IRF, time window and background level as the reconstruction. Because the emitter has only two parameters and every pixel reports on the same $\tau$, the joint Fisher information matrix is $2 \times 2$ and exactly invertible; no regularization and no free constants enter. Across the photon sweep the converged pixel-wise estimator followed its single-pixel bound at a median ratio of 1.01, the oracle followed the pooled bound at 1.02, and SPOOL sat at 1.18 times the pooled bound.

***Central-pixel photon fraction and analytic gain.***

In the main-text derivation, $h_0$ denotes the discrete fraction of an emitter's photons detected in its central pixel, not the value of an unintegrated continuous probability density. The exact background-free gain in standard deviation relative to central-pixel estimation is therefore $1/\sqrt{h_0}$. For a finely sampled, normalized two-dimensional Gaussian PSF,

$$h_0 \simeq \frac{1}{2\pi\sigma_{\mathrm{PSF}}^2}, \tag{S4}$$

which gives

$$\frac{\sigma_\theta^{\mathrm{pixel}}}{\sigma_\theta^{\mathrm{pool}}} \simeq \sigma_{\mathrm{PSF}}\sqrt{2\pi}. \tag{S5}$$

The gain is linear in $\sigma_{\mathrm{PSF}}$ because the number of detector pixels covered by the PSF scales as $\sigma_{\mathrm{PSF}}^2$, whereas the standard-deviation gain is the square root of that number. We use the exact $1/\sqrt{h_0}$ computed from the pixel-integrated PSF throughout; the two differ by less than 2% at every sampling analyzed here, and by more than 5% only at and below Nyquist sampling.

This quantity is not the inverse participation ratio $1/\sum_{\mathbf{r}} h(\mathbf{r})^2$. For a normalized two-dimensional Gaussian, the latter is $4\pi\sigma_{\mathrm{PSF}}^2$, whereas the Fisher-information ratio between whole-footprint and central-pixel estimation is $1/h_0 = 2\pi\sigma_{\mathrm{PSF}}^2$. Substituting the inverse participation ratio would therefore overstate the predicted information gain by a factor of two.

***Sampling sweep.***

To establish that the gain is a property of the sampling rather than a tuned quantity, the single-emitter analysis was repeated at a fixed 200-nm PSF over sampling densities from 1.5 to 7 pixels per FWHM, corresponding to pixel pitches from 133.3 to 28.6 nm, at both one and five photons per foreground pixel. The pixel-wise dispersion was independent of sampling to within 4%, because the central pixel receives the same photon count in every condition, whereas the joint-estimator standard deviation decreased approximately as $\sigma_{\mathrm{PSF}}^{-1.00}$. The measured gain was therefore proportional to the PSF width, with fitted slopes of 0.70 per pixel-per-FWHM at one photon and 0.91 at five photons per foreground pixel ($R^2 = 0.98$ for both), compared with the theoretical slope of 1.06 implied by $\sigma_{\mathrm{PSF}}\sqrt{2\pi}$. The residual difference is accounted for by how far each estimator lies from its respective bound rather than by the theoretical prediction. At one photon per pixel, the bounded lifetime dictionary shrinks the pixel-wise estimate below its unbiased bound, a constraint that is largely released by five photons. Meanwhile, the joint estimator remains at approximately 1.20 times the pooled bound across the examined sampling conditions and photon levels, consistent with a nearly constant reconstruction penalty relative to the pooled bound.

***Scope of the oracle reference.***

The pooled bound assumes an isolated source with known position and support and a continuous matched decay model. SPOOL instead reconstructs $K$ amplitude values at every source-space pixel using a finite dictionary, nonnegativity, and early stopping. The oracle bound is therefore used as a ruler for the information made available by the PSF, rather than as the exact bound of the practical full-field estimator.

### S4 Multi-emitter simulation and baseline implementations

***Scene generation.***

Each multi-emitter scene contained 15 point sources on a $100 \times 100$-pixel field sampled at 50 nm per pixel and imaged through the same pixel-integrated 200 nm-FWHM Gaussian PSF used for the single-emitter analysis ($\sigma_{\mathrm{PSF}} = 1.699$ px). Photon counts were generated from the spatial–contrast forward model using 256 temporal bins of 0.0977 ns and the same 150 ps IRF. The background was scaled proportionally with the foreground signal at each photon level, maintaining a mean foreground-to-background ratio of 50. This is consistent with photon thinning, which reduces signal and background counts by the same fraction and therefore preserves their relative ratio. Every reconstruction used the lifetime-agnostic $K = 11$ dictionary of Sec. S2, so no method was given the true lifetimes.

The homogeneous phantom contained 15 point emitters, all assigned the same 2 ns lifetime. It therefore provides a spatially extended test case with a known and uniform ground-truth lifetime. The phantom used the same pixel-integrated 200 nm-FWHM Gaussian PSF and 50 nm pixel sampling as the single-emitter analysis, allowing the two simulations to be compared directly.

A second, heterogeneous phantom assigned each emitter a lifetime of either 2 or 4 ns, requiring the reconstruction to distinguish different temporal signatures within overlapping PSF footprints. Both lifetimes lie exactly midway between reconstruction-dictionary nodes (2.0 between 1.8 and 2.2, 4.0 between 3.8 and 4.2), the least favorable position for grid quantization. Eight independent scenes were simulated at each of five photon budgets (50 to 800 photons per bead). Photon levels in the main figures denote the mean counts over the foreground, defined as in the experimental panels by pixels above 15% of the peak intensity, and computed from the noiseless expectation so that the axis value itself carries no shot noise. The corresponding counts at the emitter-center pixels, which are larger by a factor of about 2.1 in these scenes, are recorded alongside them in the released results files.

All methods were applied to identical photon-count cubes using the same emitter positions, random seeds, photon budgets, and forward model. Sequential unmixing–deconvolution controls were evaluated on the homogeneous phantom so that their performance reflected the order of operations rather than errors in assigning lifetime species.

Except where noted, the true PSF used to generate the data was supplied to each model-based method. The true decay bases were not: every reconstruction used the same lifetime-agnostic $K = 11$ dictionary of Sec. S2, and the true lifetimes entered scene generation only. The FPFLI baseline takes no explicit forward model as input and relied instead on weights retrained on this forward model, as described below.

##### *NC-PCA denoising + NNLS.*

We benchmarked against the noise-corrected PCA (NC-PCA) denoiser of Soltani et al. [1], following their released implementation rather than a generic truncated-SVD denoise. The defining step is a per-bin noise correction applied *before* the decomposition: each time-slice image is divided by the square root of its own spatial mean, an approximate Poisson noise scale, so that the decomposition is not dominated by whichever bins happen to carry the most absolute counts. The normalized data are then projected onto their $r$ leading principal components and reconstructed, and the same per-bin factors are multiplied back to restore the original scale. We used $r = 3$, the value the authors report as optimal in their own study, rather than tuning the rank against our known ground truth. Each denoised per-pixel curve was then unmixed against the same known bases using nonnegative least squares [2].

##### *Fit-free phasor analysis.*

A standard phasor baseline [3]: for each pixel, the phasor coordinates $G(\mathbf{r})$ and $S(\mathbf{r})$ were computed as the cosine- and sine-weighted, intensity-normalized moments of the decay at 80 MHz, and converted to a single-exponential lifetime via $\tau = S/(\omega G)$. At these photon counts $G$ can become small or negative purely from Poisson noise, so $\tau$ diverges or turns non-physical for a fraction of estimates; we report phasor RMSE over the physically-plausible subset ($0 < \tau < 10$ ns) and state the excluded fraction in the main text. The two scenes fail at different rates for a physical reason: at 80 MHz the phasor coordinate of the 4 ns species lies far closer to the origin than that of the 2 ns species, and is therefore far more easily driven negative by shot noise, so the heterogeneous scene – the one that contains it – fails more often. At the lowest photon budget the excluded fraction is 18% in the homogeneous scene and 38% in the heterogeneous scene; in the homogeneous scene it falls to 2% by the highest budget, whereas in the heterogeneous scene it remains between 22 and 40% at every budget examined.

This is an unfavorable setting for phasor analysis, and deliberately so in one respect but not in another. Practitioners working at these photon counts do not compute a raw per-pixel phasor: they bin spatially, median-filter the $G$ and $S$ maps, or pool over segmented regions, and any of these would substantially improve the numbers reported here at the cost of spatial resolution. We do not apply them because we wanted every method to see the identical per-pixel data, and because SPOOL's own gain comes precisely from using neighboring photons – so allowing one method to pool spatially while forbidding it to another would confound the comparison. The consequence, which we state rather than hide, is that the phasor curve should be read as the performance of the *unbinned* estimator, not as the best a practitioner could obtain.

##### *Retrained FPFLI baseline.*

We benchmarked against FPFLI [4], a two-stage deep-learning pipeline designed for few-photon FLIM: a local lifetime estimator (LLE), a ConvMixer-style 1D network that takes a pooled decay curve together with the system IRF and regresses a local lifetime; and a neural implicit image interpolation stage that reconstructs a full-resolution lifetime image from those coarser estimates, guided by the raw intensity image. FPFLI operates in detector space: it uses spatial context and intensity information, but does not invert the optical PSF, so it is not expected to recover deconvolved source structure. We evaluate its estimate at the same true bead locations as the other methods, on its native detector-space grid.

*Why it was retrained.* The authors' released weights were trained on a distribution that differs from this benchmark in four ways, all read directly from their released synthetic data preparation scripts: a bin width of 0.039 ns (a 9.98 ns window) against 0.0977 ns (25.0 ns) here; an IRF of 167 ps against 150 ps; a photon budget of 7–10 counts per pixel against 1.3–21 across our sweep; and, most importantly, *no optical blur at all* – their generator assigns an independent decay to each pixel of a reference image without any spatial convolution. The last is structural rather than a distribution shift, and retraining cannot remove it; retraining removes the first three, so that what remains is the architectural difference alone.

*Training data.* The LLE consumes what its own inference code feeds it, which is not a single pixel's decay: the adapter sums $8 \times 8$ spatial blocks, normalizes each summed curve by its maximum, and multiplies the network output by $\Delta t \times 100$. Training data were therefore generated in exactly that representation — $8 \times 8$ pooled, max-normalized blocks, with targets in the network's internal units — from 120,000 blocks drawn from the forward model of this work, with lifetimes uniform over 1–5 ns. Block photon counts were stratified over six bands spanning 50–8000 counts, which brackets the 83–1330 counts a block receives across the benchmark sweep; without stratification the dim tail of the PSF would place most of the training set below the range at which the network is actually asked to predict. Per-block targets are the amplitude-weighted mean lifetime of the *blurred* component maps, the quantity a pixel of a blurred image can report and the detector-space analogue of Eq. (6). Training and evaluation used disjoint random seeds.

*Training and selection.* The authors' training script and hyperparameters were used unchanged apart from a NumPy 2.0 compatibility fix. Checkpoints were selected by re-evaluating each on a held-out fifth of the training distribution rather than by the validation loss recorded during training: that script computes its validation loss from a single mini-batch, and on an earlier run it ranked three independently initialized models in the opposite order to their full-set performance. The selected network reaches an RMSE of 0.209 ns on the held-out set, falling from 0.35 ns at 50–100 counts per block to 0.09 ns above 2000. This is comparable to an approximate photon-statistical reference of 0.215 ns, computed for matched single-exponential decays by combining the per-photon Fisher information with the uniform 1–5 ns prior, which suggests that the retrained network is limited mainly by photon statistics rather than by architecture or optimization. That reference is indicative only: 92% of the pooled blocks contain both lifetime species, so a single-exponential calculation does not describe the problem the network actually solves, and the two numbers should not be read as a bound and its attainment.

*How its photon budget is obtained.* The $8 \times 8$ pooling means the LLE obtains its photon budget the same way the $3 \times 3$-binned control of Fig. 2 does — by discarding spatial sampling, here reducing the number of spatial samples by a factor of 64, equivalent to eightfold coarser sampling along each lateral axis — and recovers resolution afterwards by intensity-guided interpolation rather than by inverting the PSF. This is favorable to the strategy in the homogeneous phantom, where a block usually reports a single lifetime, and unfavorable in the heterogeneous one, where it does not. The two adapter thresholds also matter at the lowest budgets: pixels below 3 counts and blocks below 50 counts are discarded by the published method before the network is reached, which removes most of the field at 1.3 photons per foreground pixel.

***Sequential (two-stage) controls.***

SPOOL deconvolves the PSF and unmixes the dictionary inside a single Poisson likelihood. Three controls perform the same two operations *sequentially*, to test whether the joint treatment is what matters as opposed to the mere presence of a deconvolution step. Each first unmixes the measured decay at every pixel independently by nonnegative least squares against the same dictionary [2], producing detector-space amplitude maps, and then deconvolves each of those maps spatially with the same known PSF. They differ only in the deconvolution used:

- **Wiener.** Classical regularized inverse filter, $\hat{X} = \bar{H}\,Y/(|H|^2 + \mathrm{SNR}^{-1})$ with $\mathrm{SNR} = 50$, applied in the frequency domain to each amplitude map. The kernel is rolled so that its center sits at the origin and the transfer function is zero-phase; padding it as the forward convolution does would displace the deconvolved map by one kernel width.
- **Total variation.** Gradient descent on $\frac{1}{2}\|h * x - y\|_2^2 + \lambda\,\mathrm{TV}(x)$ with $\lambda = 0.02$, 100 steps, and a non-negativity projection at each step.
- **Richardson–Lucy.** The *same* multiplicative Poisson update, the *same* PSF and the *same* damping ($\eta = 0.9$) used by SPOOL, applied for 50 iterations to each amplitude map.

The first two minimize a *least-squares* data-fidelity term and therefore assume additive Gaussian noise of uniform variance. The amplitude maps they operate on are neither: at these photon counts the pixel-wise NNLS output is heteroscedastic and markedly non-Gaussian. That mismatch is part of what the comparison is designed to expose, and we state it rather than leave it implicit.

Wiener and total-variation deconvolution differ from SPOOL in three respects at once — the order of operations, the deconvolution algorithm, and the noise model — so neither can attribute the failure of the two-stage route to any one of them. The Richardson–Lucy control removes one of the three: it applies the identical Poisson update with the identical PSF and damping. It does not remove all of them, and we state this rather than claim more than the control supports. Its first stage is a least-squares unmixing, so it differs from the joint estimator both in *when* the deconvolution runs and in the noise model assumed by the preceding decomposition. A control matching the noise model at both stages – pixel-wise Poisson unmixing followed by the same Richardson–Lucy update – would isolate the ordering alone and is the natural next test. As implemented, this control recovers at most 27% of the distance between pixel-wise estimation and joint inference, and at the lowest photon budget it degrades the pixel-wise estimate rather than improving it (Fig. S1). It is nonetheless the best of the three sequential controls above two photons per foreground pixel, with Wiener marginally ahead below that and total variation worse than no deconvolution at every budget.

Hyperparameters were fixed across photon levels rather than tuned per level, matching the treatment of every other baseline. All three controls, the pixel-wise MLE and the proposed estimator were run on identical data, with the same scenes, seeds, photon budgets and 8 independent trials as the main simulation benchmark; the pixel-wise MLE and SPOOL curves reproduce those of the homogeneous panel in the main text to three decimal places.

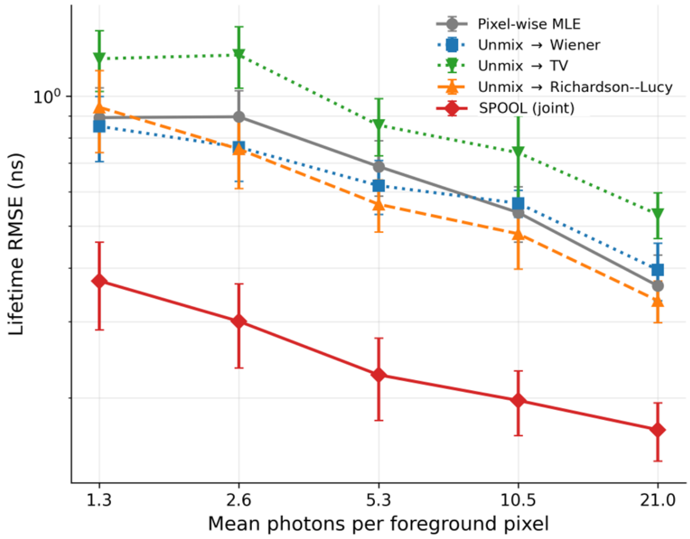


**Fig. S1 Sequential unmixing and deconvolution do not reproduce joint inference.** Lifetime RMSE is shown for the homogeneous 15-bead phantom ($\tau = 2$ ns) as a function of the mean number of detected photons per foreground pixel. Curves show mean ± s.d. across eight independent trials generated with identical scenes, seeds, photon budgets, and the same PSF and lifetime dictionary. Each sequential method first applies pixel-wise nonnegative unmixing and then deconvolves the recovered amplitude maps. The Richardson–Lucy control uses the same PSF, multiplicative Poisson update, damping ($\eta = 0.9$), and 50 iterations as the joint estimator, and differs from it both in the ordering of the two operations and in the least-squares noise model of its first stage. Joint inference achieves a 2.0–2.5-fold lower RMSE than the best sequential control at every photon budget. Richardson–Lucy is the best of the three above two photons per foreground pixel and Wiener below it. Total variation is worse than applying no deconvolution at all at every budget, and Wiener becomes so above ten photons per pixel: the pixel-wise unmixing that precedes them produces heteroscedastic, markedly non-Gaussian amplitude maps whose noise a least-squares deconvolution amplifies.

#### S5 Experimental samples and acquisition parameters

***Fluorescent beads.***

Yellow-green fluorescent beads from Invitrogen (approximately 40 nm in diameter) were imaged using a custom two-photon excitation FLIM microscope at a 30 nm pixel size with 256 time bins. Because the beads are compact and well separated, they provide the experimental counterpart of the isolated point sources used in the simulation benchmark.

***Single-labeled cells.***

Microtubules were imaged using a custom confocal FLIM microscope. Briefly, HeLa cells grown on high-precision (1.5H) glass coverslips were fixed, and microtubules were immunolabeled using an anti-tubulin primary antibody and an Oregon Green-conjugated secondary antibody. The coverslip was mounted in aqueous mounting medium and sealed at its edges with clear nail polish, and imaged at a 39 nm pixel size with 256 time bins. It provides an extended, filamentous structure in which neighboring pixels are strongly correlated. Only a single fluorophore is present, so the lifetime is expected to be approximately uniform relative to the two-species samples.

***Dual-labeled cells.***

A commercially prepared, pre-mounted two-color FLIM reference slide (GATTA-Cells, GATTAquant GmbH, Germany) contained fixed U2OS cells immunostained against TOM20, a mitochondrial outer-membrane marker, and $\alpha$-tubulin, a microtubule marker, visualized with Alexa Fluor 488 and Alexa Fluor 555, respectively, and mounted in ProLong Diamond antifade medium. Cells were simultaneously excited at 516 nm and imaged at $520 \times 520$ pixels with a 60.4 nm pixel size and 150 time bins. The lifetime varies spatially between the two labeled structures, so no single scalar ground truth exists. Foreground pixels were defined as those exceeding a fixed intensity threshold, corresponding to a mean of 435.4 photons per pixel in the full, unthinned acquisition.

***Hyperspectral sample.***

A fixed, dual-labeled huFIB cell contained an extended microtubule network visualized by Alexa Fluor 555–tubulin and a punctate mitochondrial compartment visualized by an Alexa Fluor 488 probe. The sample was acquired as a unidirectional $xy\lambda$ Lambda scan at 200 Hz (pixel dwell time 3.8 µs), with 514 nm excitation, detection from 520 to 645 nm across 26 spectral channels at 5 nm spacing, and a 50 nm pixel size. Raw integer photon counts were retained per spectral channel; no on-instrument binning, smoothing or background subtraction was applied before reconstruction. Reconstructions were performed on a $512 \times 512$ center crop of the $1024 \times 1024$ acquisition, with a full-photon mean of 215 photons per pixel over the foreground.

***Imaging setups.***

Single-emitter data were acquired in previous work using custom-made time-resolved microscopes equipped with a TCSPC card (SPC-830, Becker & Hickl). Two-photon excitation FLIM images were acquired using a femtosecond laser operating at a repetition rate of 80 MHz and a wavelength of 760 nm [5], whereas confocal FLIM images were acquired using a picosecond supercontinuum source spectrally filtered at 488 nm [6].

Multi-emitter data were acquired on a Leica STELLARIS confocal platform (Boston Innovation Hub) with an HC PL APO CS2 63×/1.40 oil-immersion objective lens and the pinhole set to 1 Airy unit. Excitation was provided by the STELLARIS supercontinuum white-light laser operating at 80 MHz, and fluorescence was detected on HyD S detectors. For the time-resolved measurements, photon arrival times were recorded with the FALCON module and stored in PTU format; decay histograms were computed from the PTU temporal stacks with custom Python code.

The experimental datasets were acquired independently using acquisition settings appropriate for each specimen and imaging modality, resulting in different scanning magnifications and native sampling pitches. The four datasets are therefore analyzed at their native sampling: 30 nm for the beads, 39 nm for the microtubules, 50 nm for the hyperspectral cell, and 60.4 nm for the dual-labeled FLIM cell. Given the 200-nm PSF full width at half maximum, these pitches correspond to 6.7, 5.1, 4.0, and 3.3 detector pixels per PSF full width at half maximum, respectively. Because the predicted gain depends on this sampling ratio, each dataset is evaluated against its corresponding theoretical prediction rather than against a single common value.

#### S6 Photon-thinning and evaluation protocol

***Thinning.***

No ground-truth parameter map exists for a real sample. Photon-budget sweeps were therefore generated from each full-photon acquisition by binomial thinning of the raw photon-count cube: each recorded photon is retained independently with probability $p = n/n_{\mathrm{full}}$, where $n$ is the target mean photon count per foreground pixel. Because a Poisson process thinned binomially is again Poisson, the thinned cube has exactly the statistics of a shorter acquisition of the same specimen. Six independently generated thinning realizations were produced at each photon level (eight independent scenes for the simulation benchmark), and the background term was scaled by the same $p$. These replicates quantify thinning variability conditional on the acquired full-photon dataset; they are not independent specimen acquisitions, and the reported error bars should be read accordingly. Metrics were computed by pooling all replicates and all mask pixels rather than by averaging per-replicate values.

***Photon budgets.***

All photon budgets in this work, simulated and experimental, are quoted as the mean number of detected photons over the foreground, defined as the pixels receiving at least 15% of the peak intensity. This single rule makes the horizontal axes of the simulation and experimental figures directly comparable. For an isolated emitter the ratio between the central pixel and this foreground mean is fixed by the PSF geometry at 2.3.

**Table S1** Reconstruction parameters for the FLIM datasets. The same estimator is used throughout; only the forward-model inputs differ. The PSF is computed from the objective (FWHM = $0.51\lambda/\mathrm{NA}$, NA = 1.40, $\lambda = 550$ nm $\Rightarrow$ 200 nm) rather than fitted, so it is fixed by the optics rather than tuned, and integrated over each detector pixel. The bead column is the exception: that dataset was acquired by two-photon excitation at 760 nm and its 200 nm width follows from $0.51\lambda_{\mathrm{ex}}/(\mathrm{NA}\sqrt{2}) = 196$ nm instead (Sec. S7). The predicted gain is the exact $1/\sqrt{h_0}$ of the pixel-integrated PSF; the finely-sampled approximation $\sigma_{\mathrm{PSF}}\sqrt{2\pi}$ is smaller by less than 2% in every column. The decay dictionary is lifetime-agnostic: no dye lifetime enters the model.

| | **Multi-emitter simulation** | **Beads** | **Microtubules** | **Cells** |
|---|---|---|---|---|
| *Acquisition* | | | | |
| Pixel size (nm) | 50 | 30 | 39 | 60.4 |
| Time bins | 256 | 256 | 256 | 150 |
| Bin width $\Delta t$ (ns) | 0.0977 | 0.0977 | 0.0977 | 0.0970 |
| Field of view (px) | $100^2$ | $100^2$ | $200^2$ | $520^2$ |
| *Optics* | | | | |
| Objective NA | — | 1.40 | 1.40 | 1.40 |
| PSF FWHM (nm) | 200 | 200 | 200 | 200 |
| PSF $\sigma_{\mathrm{PSF}}$ (px) | 1.70 | 2.83 | 2.18 | 1.41 |
| Pixels per PSF FWHM | 4.0 | 6.7 | 5.1 | 3.3 |
| Predicted gain $1/\sqrt{h_0}$ | 4.3× | 7.1× | 5.5× | 3.6× |
| *Instrument response* (Gaussian, reconvolution fit to the summed decay) | | | | |
| $t_0$ (ns) | 0.500 | 12.265 | 11.830 | 0.821 |
| $\sigma_{\mathrm{IRF}}$ (ns) | 0.064 | 0.049 | 0.064 | 0.067 |
| IRF FWHM (ps) | 150 | 116 | 150 | 157 |
| *Background* | | | | |
| Counts per pixel per bin | † | estimated per dataset, Eq. (S2) | | |
| *Decay dictionary* (lifetime-agnostic; no dye lifetime assumed) | | | | |
| $K$ | 11 | 11 | 11 | 11 |
| $\tau$ range (ns) | 1.0–5.0 | 1.0–5.0 | 1.0–5.0 | 1.0–5.0 |
| $\tau$ spacing (ns) | 0.4 | 0.4 | 0.4 | 0.4 |
| True $\tau$ (generation only) | 2.0, 4.0 | — | — | — |
| *Reconstruction* | | | | |
| Iterations | 50 | 50 | 50 | 50 |
| Damping $\eta$ | 0.9 | 0.9 | 0.9 | 0.9 |
| Trials / thinning replicates | 8 | 6 | 6 | 6 |
| Amplitude mask (% of own max) | — | 5 | 5 | 5 |

†Scaled proportionally with the foreground signal at each photon level to maintain a mean foreground-to-background ratio of 50; the single-emitter study instead uses a fixed $10^{-4}$ counts per pixel per bin (Sec. S1), which is a ratio of 39 at one photon per foreground pixel. Sec. S8 sweeps this ratio from 50 to 5. The true lifetimes enter scene generation only: every reconstruction uses the lifetime-agnostic $K = 11$ dictionary, and 2.0 and 4.0 ns lie midway between its nodes.

**Table S2** Hyperspectral experiment. The estimator, the PSF model and every numerical setting are unchanged from the FLIM experiments; only the dictionary differs – Gaussian emission bands in place of IRF-convolved exponential decays.

| | |
|---|---|
| **Specimen** | Fixed huFIB cell, dual-labeled: Alexa Fluor 555–tubulin (extended microtubule network) and an Alexa Fluor 488 mitochondrial probe (punctate) |
| **Acquisition** | STELLARIS confocal, HC PL APO CS2 63×/1.40 oil, 1 AU pinhole |
| Excitation | 514 nm |
| Detection band | 520–645 nm |
| Spectral channels | 26 (5 nm spacing) |
| Pixel size | 50 nm |
| Scan | unidirectional $xy\lambda$ Lambda scan, 200 Hz (3.8 µs dwell) |
| Field analyzed | 512 × 512 center crop of the 1024 × 1024 acquisition |
| Full-photon mean | 215 photons per foreground pixel |
| **Optics** | FWHM = $0.51\lambda/\mathrm{NA} = 0.51 \times 550/1.40 = 200$ nm |
| PSF $\sigma_{\mathrm{PSF}}$ | 1.70 px (4.0 pixels per FWHM) |
| Predicted gain | $1/\sqrt{h_0} = 4.3\times$ |
| **Spectral dictionary** | $K = 11$ Gaussian emission bands |
| Centers | 532–632 nm, 10 nm spacing |
| Width | 40 nm FWHM each, unit-area normalized |
| Source | Constructed to tile the detection band. *No fluorophore spectrum is used*: the dictionary is the direct analogue of the lifetime-agnostic decay dictionary, and the dyes' published emission spectra enter neither the model nor the fit. |
| **Reconstruction** | 50 iterations, $\eta = 0.9$, 6 thinning replicates |
| Amplitude mask | 3% of each method's own maximum |
| Recovered quantity | amplitude-weighted mean emission wavelength, $\hat{\lambda}_{\mathrm{src}}(\mathbf{r}) = \sum_k \lambda_k A_k(\mathbf{r})/[\sum_k A_k(\mathbf{r}) + \epsilon]$ for SPOOL; the same expression uses detector-space amplitudes $\tilde{A}_k$ for the pixel-wise baseline |

***Reference-relative metrics.***

Each method is compared against *its own* full-photon reconstruction, computed at the same iteration count as the thinned reconstructions. This is essential and easy to get wrong: reusing a stored high-photon reference produced at a different iteration count would compare each method against a differently-converged version of itself. The three quantities reported are, for a method $M$ with thinned estimate $\hat{\theta}_M^{(n)}$ and full-photon reference $\hat{\theta}_M^{\mathrm{full}}$, over the pixels in that method's own mask $\mathcal{M}_M$:

$$\text{mean signed reference error} = \big\langle \hat{\theta}_M^{(n)} - \hat{\theta}_M^{\mathrm{full}} \big\rangle_{\mathcal{M}_M}, \tag{S6}$$

$$\text{reference-relative s.d.} = \mathrm{sd}\big(\hat{\theta}_M^{(n)} - \hat{\theta}_M^{\mathrm{full}}\big)_{\mathcal{M}_M}, \tag{S7}$$

$$\text{reference-relative RMSE} = \left\langle \left(\hat{\theta}_M^{(n)} - \hat{\theta}_M^{\text{full}}\right)^2 \right\rangle_{\mathcal{M}_M}^{1/2}. \tag{S8}$$

The RMSE is defined directly and satisfies $\text{RMSE}^2 = \text{bias}^2 + \text{s.d.}^2$ when all three are evaluated over the same pooled set. These are repeatability metrics, not errors against a truth, and we do not interpret them as absolute accuracy or as Cramér–Rao efficiency. One consequence is used explicitly in Sec. S8: because the reference is produced by the same method under the same assumed model, it inherits any error in that model, and the reference-relative metrics are therefore insensitive to model mismatch by construction. The sensitivity analysis is scored against the simulated ground truth for that reason.

***Masks.***

A pixel enters a method's statistics, and appears in its displayed map, only where that method's own recovered amplitude exceeds a fixed fraction of its own maximum: 5% for the FLIM samples, 3% for the hyperspectral sample. A pixel at which a method reconstructed no amplitude carries no information, and including it would score an estimate that was never made. The two methods are consequently evaluated on different pixel counts because support selection can influence the experimental gain ratios.

***Which dispersion is plotted.***

For the single-species samples the mean signed reference error is small at every photon level, so the reference-relative standard deviation and RMSE coincide to within a few percent. The mean recovered lifetime is plotted with the standard deviation shown as error bars. For the two-species samples the signed term is not negligible – the pixel-wise estimator approaches its own high-photon limit from a systematic offset – so RMSE is plotted, which reports the full departure. Both signed and dispersion components are reported for the two-species datasets.

***Power-law fits.***

Photon-scaling exponents were estimated by ordinary least squares after logarithmic transformation,

$$\log e(n) = \alpha \log n + c, \tag{S9}$$

where $e(n)$ is the reference-relative error and $n$ is the mean photon count per foreground pixel. Fits used all photon levels in the sweep. Because the photon levels were generated by binomial thinning of the same full-photon acquisition, measurements across photon levels are correlated. Intervals obtained from ordinary least squares therefore underestimate the total uncertainty and are reported as descriptive intervals rather than as formal tests of estimator efficiency.

### S7 PSF specification and iteration count

***PSF specification.***

A sample-specific PSF measurement was not available, so one nominal diffraction-limited lateral width was used throughout. For the 63×/1.40-NA oil-immersion objective, the PSF full width at half maximum was calculated as

$$\mathrm{FWHM} = 0.51\lambda/\mathrm{NA}, \tag{S10}$$

using a representative emission wavelength of $\lambda = 550$ nm. This gives FWHM = 200 nm and $\sigma_{\mathrm{PSF}} = 85$ nm for the Gaussian approximation.

The same physical width was converted to detector pixels using each dataset's sampling: $\sigma_{\mathrm{PSF}} = 2.83$ pixels for beads, 2.18 pixels for microtubules, 1.41 pixels for dual-labeled FLIM cells, and 1.70 pixels for the hyperspectral sample. The baseline single- and multi-emitter simulations were sampled at 50 nm and used $\sigma_{\mathrm{PSF}} = 1.70$ pixels. In the sampling-sweep analysis, the physical PSF width was held fixed at 200 nm while the pixel pitch was varied from 133.3 to 28.6 nm, corresponding to 1.5–7 pixels per PSF full width at half maximum.

The bead dataset requires a separate justification, because it was acquired by two-photon excitation at 760 nm rather than by one-photon confocal imaging, so the emission-based rule above does not apply to it. For two-photon excitation the signal follows the square of the excitation intensity, which narrows a Gaussian excitation focus by $\sqrt{2}$, giving $0.51\lambda_{\mathrm{ex}}/(\mathrm{NA}\sqrt{2}) = 0.51 \times 760/(1.40\sqrt{2}) = 196$ nm. The two derivations are physically unrelated but agree to within 2%, so the same 200 nm width is used for that dataset as well; the agreement is a coincidence of these particular wavelengths and should not be taken as a general rule.

A single, spatially invariant and wavelength-independent PSF was used for every reconstruction. The relevant one-photon fluorophores span approximately 515–565 nm in peak emission, over which the nominal diffraction-limited width varies by less than 10%, from 188 to 206 nm. The PSF width was fixed before reconstruction and was not fitted to the measured performance. The assumed width enters both the reconstruction and the optics-derived prediction, so agreement between predicted and measured gains is a consistency check rather than an independent validation of PSF accuracy. Sec. S8 therefore reconstructs the single-emitter data with deliberately incorrect PSF widths and reports how far the measured gain moves. Joint estimation of the PSF alongside the amplitude maps, which would remove the assumption rather than bound its consequences, is left to future work.

###### ***Choice of iteration count.***

Multiplicative Richardson–Lucy-type updates are semi-convergent: additional iterations initially reduce deterministic reconstruction bias, but continued iteration progressively amplifies shot noise and can increase the total estimation error [7]. We therefore treated the iteration count as a fixed bias–variance operating point rather than a convergence criterion, and selected it from a ground-truth single-emitter simulation instead of tuning it separately for each dataset.

The scan used the nominal 200 nm-FWHM PSF and photon levels spanning 1–15 detected photons per foreground pixel. The iteration count giving the smallest lifetime RMSE depends on photon level: at the lowest counts the minimum occurs earlier, whereas higher-count data tolerate and can benefit from additional iterations (Fig. S2a). A single per-dataset optimum would therefore amount to photon-level-specific tuning and would not provide a common reconstruction setting for the experimental photon sweeps.

We instead averaged the lifetime RMSE over the complete 1–15 photons-per-pixel range and examined that range-averaged error as a function of iteration count (Fig. S2b). For the nominal PSF, the curve reaches a broad, shallow minimum in the 50–75-iteration range. The same analysis was repeated after deliberately perturbing the PSF width supplied to the reconstruction by $\pm 10\%$, approximately bracketing the wavelength-dependent variation expected over the relevant emission band. The minima remain in the same 50–75-iteration range, with 50 iterations lying near the minimum for all three PSF assumptions. We therefore fixed the reconstruction at 50 iterations and $\eta = 0.9$ for every simulated and experimental dataset, including the hyperspectral reconstruction. This choice is not claimed to be a universal optimum; it is a single robust operating point chosen before comparison so that no dataset or photon level receives its own tuned stopping rule.

The damping exponent $\eta = 0.9$ was likewise held fixed throughout and was not re-optimized for individual datasets. To test sensitivity to this choice, we swept $\eta$ from 0.7 to 1.0 while otherwise keeping the reconstruction conditions unchanged. Across this range, the reconstruction error varied by less than 1%, indicating that the reported performance is not sensitive to the specific damping value used. On the experimental single-species samples, later iterations also produced progressively more concentrated source-space amplitude maps and weaker agreement with the corresponding detector-space summed-intensity image. Because a deconvolved source-space image is expected to be sharper than that detector-space projection, this observation is treated only as a qualitative consistency check; the ground-truth RMSE analysis above is the basis for selecting the iteration count.

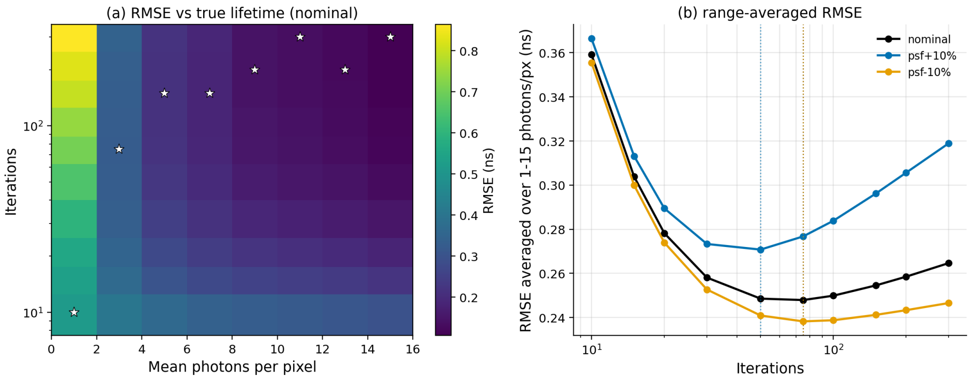


**Fig. S2 Selection of a fixed reconstruction iteration count.** **a**, Lifetime RMSE relative to the known 2 ns lifetime in the single-emitter simulation as a function of mean detected photons per foreground pixel and iteration count, using the nominal 200 nm-FWHM PSF. Stars mark the iteration count giving the lowest RMSE at each photon level. The optimal stopping point shifts to later iterations as the photon budget increases, showing that no single iteration count is pointwise optimal across the full photon range. **b**, Lifetime RMSE averaged over 1–15 photons per foreground pixel as a function of iteration count for the nominal PSF and for reconstruction PSF widths perturbed by +10% and −10%. The curves exhibit broad minima in the 50–75-iteration range. A fixed value of 50 iterations lies near the minimum for all three PSF assumptions and was therefore used for every reconstruction in this work.

### S8 Sensitivity to model mismatch

The estimator is handed a PSF and a background level rather than estimating them, so the results depend on both being approximately right. This section quantifies how approximately. Three quantities are perturbed independently about their nominal values, one at a time:

- the foreground signal-to-background ratio, at 5, 10, 20 and 50;
- the assumed PSF full width at half maximum, in error by −20, −10, 0, +10 and +20%;
- the assumed background rate, in error by −20, 0 and +20%.

The geometry, dictionary, IRF, estimators and readout are those of the single-emitter study of Sec. S3, unchanged. Only the values *supplied to the estimators* are perturbed: the photon counts are always generated with the true PSF and the true background, so a mismatch degrades the estimator without altering the information content of the data. The Cramér–Rao bounds are likewise computed from the true PSF and true background and are therefore invariant across the PSF sweep by construction, which is what allows any movement in the measured dispersion to be read as lost estimator efficiency rather than lost information.

Results are quoted at one detected photon per foreground pixel, the most demanding point of the sweep, from 480 Poisson realizations per condition split into eight blocks; tabulated uncertainties are standard errors over those blocks. Common random numbers are used across conditions wherever the true model is unchanged, which is the case throughout the PSF and background sweeps, so those comparisons are exactly paired: the pixel-wise estimator, which never uses the PSF, returns bit-identical values across the whole PSF sweep. The background sweep holds the signal-to-background ratio at 10 so that a misestimated background has something to act on; at 50 the background is small enough that a 20% error in it is inconsequential and a robustness claim made there would be vacuous.

The signal-to-background ratio deserves a word, because the two simulations in this work set it differently (Sec. S1). The multi-emitter benchmark holds it at 50. The single-emitter study instead fixes the background at $10^{-4}$ counts per pixel per bin, which over the 256-bin window is 0.0256 photons per pixel and therefore a ratio of 39 at one photon per foreground pixel. The sweep below spans 5 to 50 and so brackets both, and extends to a background nearly eight times larger than either.

***Signal-to-background ratio.***

The measured gain varies by 7.1% as the ratio falls from 50 to 5 (Table S3). Most of that is not the estimator degrading. Raising the background raises both Cramér–Rao bounds and lowers the ratio between them, so the gain that is attainable at all falls from 4.10 to 3.87 over the same range; expressed as a fraction of that attainable gain, the measured value varies by 2.0%. SPOOL sits between 1.13 and 1.17 times the pooled bound in standard deviation at every ratio tested, so its efficiency is essentially independent of the background level.

The bias behaves differently and is worth stating explicitly: it rises from +0.119 ns at a ratio of 50 to +0.226 ns at 5. The ratio-50 condition is close to the single-emitter study's fixed-background condition, which corresponds to a ratio of 39, providing a nearby consistency check between the two analyses.

One trend runs against the naive expectation and is reported rather than suppressed. The estimator's efficiency relative to the oracle *improves* as the background rises, from 0.83 at a ratio of 50 to 0.95 at 5: the oracle, which pools 81 pixels, gains more from a clean background than SPOOL does, so the gap between them closes as conditions worsen. Whatever limits SPOOL at high signal-to-background is therefore not photon statistics.

***Background misspecification.***

A ±20% error in the assumed background changes the measured gain by 2.2% and leaves the estimator between 1.10 and 1.16 times the pooled bound. The effect on the bias is larger and signed: underestimating the background inflates the bias to +0.207 ns and overestimating it deflates it to +0.143 ns, from +0.173 ns when the background is correct. Unmodeled background is absorbed into the recovered amplitudes and deconvolved into source space along with the signal, so the practical recommendation is to estimate the background conservatively with respect to this particular bias; Eq. (S2), which averages a signal-free corner of the field, overestimates the background when faint structure is present in that corner, which reduces the lifetime bias but also suppresses the recovered amplitudes.

***PSF misspecification.***

An assumed PSF too narrow by 20% costs 13.7% of the gain; too wide by 20%, the measured gain *rises* by 5.8%. The estimator does not fall to the pixel-wise baseline anywhere in the range tested, and retains at least 86% of its matched-model gain throughout.

The asymmetry is not an artifact and resolves once the bias is examined alongside the dispersion. As the assumed PSF widens, the standard deviation falls monotonically from 1.31 to 1.07 times the pooled bound, while the bias rises monotonically from +0.159 to +0.200 ns. An over-wide assumed PSF over-deconvolves, concentrating the recovered amplitude into fewer source pixels; that suppresses the variance at the emitter pixel and is paid for in bias. At one photon per pixel the estimator is variance-limited, so the RMSE still improves slightly across the range, from 0.501 to 0.434 ns. This should not be read as a recommendation to over-specify the PSF: the variance term shrinks as the photon count rises while the bias term does not, and the gain reported here is a standard-deviation ratio in which the bias does not appear at all.

The ±20% range exceeds the wavelength-dependent variation predicted by the nominal diffraction-limited model, over which the width varies by less than 10% across the relevant 515–565 nm emission band (Sec. S7), and brackets a substantial range of additional optical mismatch. It should not, however, be read as bounding every aberration- or instrument-dependent PSF error: the effective confocal response also depends on the excitation wavelength, the coupling of excitation and detection, the pinhole setting, refractive-index mismatch and alignment, none of which the nominal model captures.

**Table S3** Sensitivity of the single-emitter reconstruction to model mismatch, at one detected photon per foreground pixel. Gain is the ratio of the pixel-wise to SPOOL's standard deviation, as in Eq. (10); the attainable gain is the corresponding ratio of Cramér–Rao bounds at the same condition. Errors are quoted against the true lifetime over 480 realizations. Bold rows are the nominal condition of each sweep. The pixel-wise estimator does not use the PSF and returns identical values throughout the PSF sweep.

| | Gain | Attainable gain | Gain / attainable | Bias (ns) | s.d. / $\mathrm{CRB}_{\mathrm{pool}}$ |
|---|---|---|---|---|---|
| *Signal-to-background ratio* (PSF and background correct) | | | | | |
| 5 | 2.85 | 3.87 | 0.735 | +0.226 | 1.160 |
| 10 | 3.01 | 3.96 | 0.760 | +0.173 | 1.128 |
| 20 | 3.07 | 4.03 | 0.763 | +0.146 | 1.141 |
| **50** | **3.07** | **4.10** | **0.748** | **+0.119** | **1.170** |
| *Assumed PSF FWHM error* (ratio 10, background correct) | | | | | |
| −20% | 2.59 | 3.96 | 0.656 | +0.159 | 1.308 |
| −10% | 2.84 | 3.96 | 0.717 | +0.164 | 1.197 |
| **0** | **3.01** | **3.96** | **0.760** | **+0.173** | **1.128** |
| +10% | 3.12 | 3.96 | 0.788 | +0.186 | 1.088 |
| +20% | 3.18 | 3.96 | 0.805 | +0.200 | 1.066 |
| *Assumed background error* (ratio 10, PSF correct) | | | | | |
| −20% | 2.95 | 3.96 | 0.745 | +0.207 | 1.160 |
| **0** | **3.01** | **3.96** | **0.760** | **+0.173** | **1.128** |
| +20% | 3.07 | 3.96 | 0.777 | +0.143 | 1.102 |

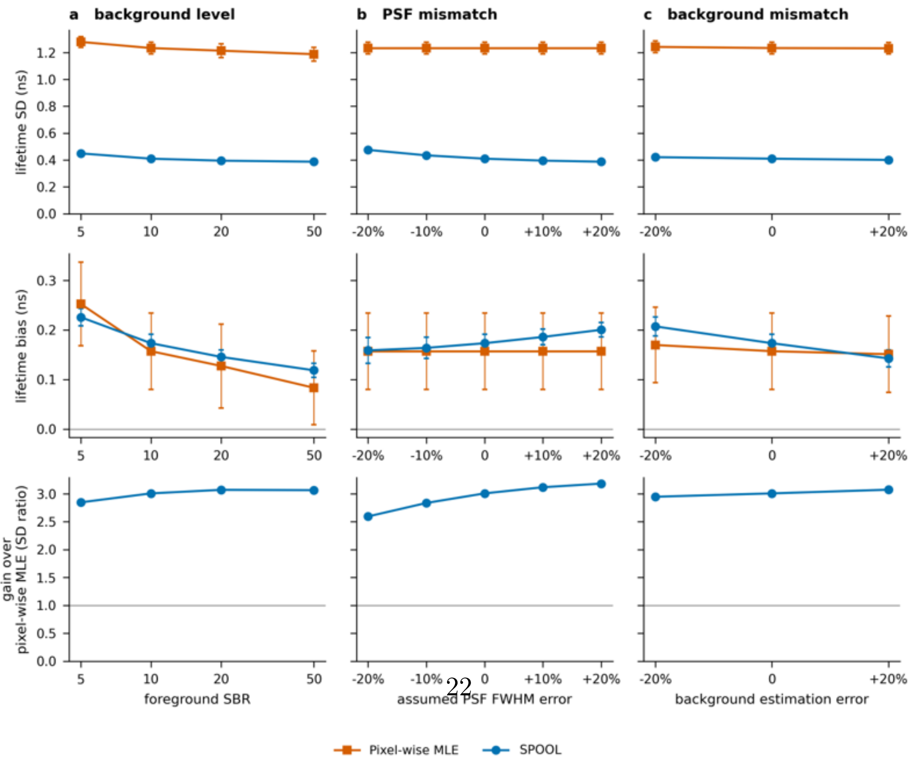




**Fig. S3 Sensitivity to model mismatch at one photon per foreground pixel.** Single-emitter Monte-Carlo simulations with the geometry, dictionary and IRF of Sec. S3 (200 nm PSF FWHM, $\sigma_{\mathrm{PSF}} = 1.70$ px, $\tau_0 = 2.0$ ns), varying the foreground signal-to-background ratio (**a**), the assumed PSF full width at half maximum (**b**) and the assumed background rate (**c**). Panels **b** and **c** hold the signal-to-background ratio at 10, low enough that a misestimated background has an effect to measure. Photon counts are always generated with the true PSF and true background; only the values supplied to the estimators are perturbed. Top row, lifetime standard deviation across 480 realizations; middle row, bias; bottom row, gain over the pixel-wise Poisson MLE, defined as the ratio of standard deviations as in Eq. (10). Error bars are standard errors over eight blocks of realizations. Common random numbers are used across conditions in **b** and **c**, where the true model is identical, so those panels are exactly paired comparisons; the pixel-wise estimator does not use the PSF and is invariant across **b** by construction. As the assumed PSF widens, the standard deviation falls and the gain rises, but the bias rises with it, from 0.159 to 0.200 ns: at this photon level the estimator is variance-limited, so the apparent benefit of an over-wide PSF is a variance–bias trade rather than an improvement in accuracy. The gray line in the bottom row marks parity with the pixel-wise estimator, which is the reference of the ratio and is therefore not plotted separately.

### S9 Runtime and computational resources

Reconstructions were implemented in PyTorch and run on an Intel Core i5-13600K CPU (14 physical cores) and an NVIDIA GeForce RTX 3060 Ti, with an equivalent NumPy CPU path available. In double precision, the CPU and GPU implementations agree to within $10^{-14}$ relative error, confirming that the GPU implementation is an optimization of the same numerical procedure rather than a separate method. The experimental reconstructions and the multi-emitter benchmark were run in double precision. The single-emitter Monte-Carlo study, which requires 400 independent realizations at each photon level, was run in single precision with TF32 matrix multiplication and verified against a double-precision run; the dictionary profiles and all Cramér–Rao bounds were evaluated in double precision throughout. The iteration-selection and sensitivity studies of Secs. S7 and S8 use the same single-precision path; the sensitivity study uses 480 realizations per condition.

All reported SPOOL reconstructions use the fixed 50-iteration setting of Sec. S7 with $K = 11$ basis components. Table S4 reports wall-clock reconstruction times for the four experimental datasets. Even in double precision, every dataset reconstructed in less than 2 s on the RTX 3060 Ti. Single-precision execution reduced the corresponding times to 0.028–0.342 s. Peak GPU memory remained below 1.9 GiB for all datasets, including the $520 \times 520 \times 150$ dual-labeled FLIM dataset. For a fixed dataset and dictionary, the computational cost per iteration is approximately constant, so wall-clock time scales approximately linearly with the iteration count.

**Table S4 Computational performance at 50 iterations and $K = 11$.** Wall-clock reconstruction times are reported for double-precision GPU (GPU f64), single-precision GPU (GPU f32), and double-precision CPU (CPU f64) execution. Peak GPU memory is the maximum memory used during reconstruction.

| | | Reconstruction time (s) | | | Peak GPU |
|---|---|---|---|---|---|
| **Dataset** | **Shape (px)** | **GPU f64** | **GPU f32** | **CPU f64** | **(GiB)** |
| Beads | $100^2 \times 256$ | 0.138 | 0.028 | 0.694 | 0.38 |
| Microtubules | $200^2 \times 256$ | 0.479 | 0.080 | 3.539 | 0.67 |
| Dual-labeled cells | $520^2 \times 150$ | 1.971 | 0.342 | 13.267 | 1.88 |
| Hyperspectral | $512^2 \times 26$ | 1.707 | 0.171 | 11.358 | 0.66 |

The dominant memory cost is the photon-ratio tensor $R = Y/(\Lambda + \epsilon)$, whose size is set by the measured photon-count cube rather than directly by the number of dictionary components. For the dual-labeled cell dataset, this tensor occupies approximately 309 MiB in double precision, compared with approximately 23 MiB for the $K = 11$ amplitude maps. Larger datasets can therefore be accommodated by processing the contrast axis in blocks, which leaves the reconstruction mathematically unchanged while reducing the peak memory required for contrast-axis intermediate tensors. No such streaming was required for any dataset analyzed here.
